# Transdifferentiation of Human Dental Pulp Mesenchymal Stem Cells into Spiral Ganglion-like Neurons

**DOI:** 10.1101/2024.02.02.578615

**Authors:** Yassine Messat, Marta Martin-Fernandez, Said Assou, Keshi Chung, Frederic Guérin, Csilla Gergely, Frederic Cuisinier, Azel Zine

## Abstract

Spiral ganglion neurons (SGN) carry auditory information from sensory hair cells (HCs) to the brain. These auditory neurons, which are the target neurons of cochlear implants, degenerate following sensorineural hearing loss (SNHL). Prosthetic devices such as cochlear implants function by bypassing lost HCs and stimulating the residual SGNs, allowing restoration of hearing in deaf patients. Emerging cell-replacement therapies for SNHL include replacing damaged SGNs using stem cell-derived otic neuronal progenitors (ONPs). However, the availability of renewable, accessible, and patient-matched sources of human stem cells constitutes a major prerequisite towards cell replacement for auditory nerve recovery. Human dental pulp stem cells (hDPSCs) extracted from human wisdom teeth are self-renewing stem cells that originate from the neural crest during development.

In this study, we developed a stepwise *in vitro* guidance procedure to differentiate hDPSCs into ONPs and then to SGNs. The procedure relies on the modulation of BMP and TGF-β pathways for neurosphere formation as a first step, then a differentiation step based on two culture paradigms exploiting major signaling pathways (Wnt, Shh, RA) and neurotrophic factors involved in early otic neurogenesis.

Gene and protein expression analyses revealed efficient induction of a comprehensive panel of known ONP and SGN-like cell markers over the course of *in vitro* differentiation. The use of atomic force microscopy revealed that hDPSC-derived SGN-like cells exhibit similar nanomechanical properties compared to their *in vivo* SGN counterparts. Furthermore, neurites extended between hDPSC-derived ONPs and rat SGN explants 4-6 days after co-culturing, suggesting the formation of neuronal contacts. These data indicate that the *in vitro* differentiated cells closely replicate the phenotypic and nanomechanical characteristics of human SGNs, advancing our culture differentiation system to the level to be used in next-generation cochlear implants and/or inner ear cell-based strategies for SNHL.

## Introduction

Sensorineural hearing loss (SNHL) is the most common type of hearing loss. Of all the types of SNHL, auditory neuropathy spectrum disorder is of particular concern. This condition, defined primarily by damage to the spiral ganglion neurons (SGNs) that relay sound transmission from the sensory organ (cochlea) to the brain with relative preservation of the hair cells (HCs), is responsible for significant hearing impairment. While the deficit of HCs can be functionally overcome by a cochlear implant, no treatment is currently available for SGN loss. Without neurons, most of the currently available cochlear implants will not function (1).

A potential therapeutic approach to the loss of sensory neurons would be to replace them by transplantation of exogenous, *in vitro* maintained, stem cell-derived otic neuronal progenitors (ONPs). These transplantations could also provide a means of delivering supportive neurotrophic factors to promote further the survival of neurons. Moreover, the delivery of ONPs at the time of cochlear implantation could extend the applicability and success rate of the current cochlear implant approach. Although part of the efforts to develop a stem cell-based therapy for neurosensory deafness aims to restore HCs, the replacement of SGNs would appear more feasible in the initial stages.

The proof of concept has been established in a previous study (2) showing that human embryonic stem cells can produce ONPs, and that these cells can partially repair a damaged cochlear nerve *in vivo*. However, for developing a cell therapy to replace SGNs, the *in vitro* generation of appropriate ONPs from reliable and easily accessible, less controversial sources of human stem cells is among the requirements for a potential clinical application.

Mesenchymal stem cells (MSCs) possess several unique properties that make them a particularly attractive source of cells for cell-based cell therapy. They have the potential to self-renew and differentiate into multiple cell types (3,4), and have been used to replace damaged cells in the nervous system using animal models of neurological disorders (5). Several lines of evidence suggest that MSCs can be used to replace damaged neurons and also to promote endogenous neuronal cell repair/survival by releasing neurotrophic factors (6,7). Furthermore, there is a rising interest in the application of MSCs to treat SNHL (reviewed in (8,9)). Some works investigated the effects of directly transplanted MSCs in the inner ear (10–12), while other studies focused on the hypothetical integration of neuro-induced MSCs (13,14).

Dental pulp-derived human mesenchymal stem cells (hDPSCs) could be an alternative source of cells as they possess both mesenchymal and neural features due to their ectodermal origin (7,15). hDPSCs could have additional advantages when compared to adult stem cells from other sources, because dental pulp epithelium and cochleo-vestibular ganglion have relatively close embryonic development and share some transcriptional pathways with the neural crest (16). Furthermore, hDPSCs are harvested from a minimally invasive, accessible location, as tooth tissues are easy to acquire via biological waste at high amounts from avulsions of wisdom teeth or premolars for orthodontic reasons (17). All these characteristics promote hDPSCs as appropriate donor cells for a potential cell therapy for SNHL and/or to provide replacement otic neurons to improve the benefits of cochlear implantation.

In the *in vivo* situation, the neurosensory cells of the inner ear are derived from the otic vesicle (18). The otic vesicle derives from non-neural ectoderm (NNE), which is induced from the ectoderm layer by a lateral-to-medial gradient of bone morphogenetic protein (BMP) signaling (19). Its ventral region contains otic sensory neuronal progenitors that give rise to primary sensory neurons, including SGNs (reviewed in (20,21)). Previous studies demonstrated that Sonic hedgehog (Shh) and retinoic acid (RA) synergistically promote the expression of sensory neuron markers and facilitate otic sensory neuronal differentiation (22). It was also shown that Shh and BMP signaling and supplementation of neurotrophic factors (i.e., brain-derived neurotrophic factor (BDNF) and NT3) play an important role in the generation of SGN-like neurons from pluripotent stem cells (2,23–25). While there is a rising interest in the use of MSCs-like human bone MSCs to generate SGN-like neurons for the appropriate properties that they offer (26,27), no *in vitro* differentiation of SGN-like neurons from adult hDPSCs through the early otic lineage has been yet proposed.

Extending on these findings and protocols, we have further assessed the possibility of deriving otic neurons *in vitro* from adult hDPSCs. We utilised two paradigms by either modulation of Wnt/Shh/RA pathways or exposure to neurotrophic factors to assess which protocol can promote otic neuronal fate specification from hDPSCs after neural induction. We then characterized cells from each paradigm at the cellular and molecular levels and added biomechanical characterization of terminally differentiated SGN-like cells. Finally, we explored the potential of the generated ONPs in a co-culture system with SGN explants for a future perspective of application in cell therapy.

We show here the stepwise *in vitro* generation of high number of human SGN-like cells from hDPSCs expressing key otic neuronal lineage gene markers and displaying the typical bipolar morphology of SGNs with the same nanomechanical properties.

## Materials and Methods

### Collection of human dental pulp and culture

The human dental pup stem cells (hDPSCs) were isolated from extracted wisdom teeth from young healthy patients (14-21 years old). Informed consent was obtained from the patients after receiving approval by the local ethics committee (Comité de protection des Personnes, Centre Hospitalier de Montpellier). We used a previously established protocol to recover pulp cells (17). Briefly, the teeth were cleaned with 2% chlorhexidine, then cut at the cementum-enamel junction by using a sterilized drill. The teeth were broken into two pieces with a scalpel and the pulp was recovered from its cavity by using a tweezers. Pulps were first cut into small pieces then digested in 2 ml solution of 3 mg/ml collagenase type 1 and 4 mg/ml of dispase (Corning). The digestion lasted 1 hr at 37°C. The cell suspension was filtered using a 70 µm strainer (Falcon) and transferred to T75 flask (Falcon) containing 10 ml of complete medium: A-MEM (Gibco), 10% FBS (Sigma), 100 µg/ml streptomycin (Sigma) and 100U/ml penicillin (Sigma). Medium was first changed after 24hr then every 3 days for 1 week. At confluency, the cells were passaged by washing the culture with DPBS (Cytiva), then detached using Trypsin-EDTA (Gibco).

### Flow cytometry

The immunophenotypic detection of mesenchymal stem cell markers i.e. (CD90, CD73, CD105) was performed by flow cytometry. Dental pulp cells at passage 4 were rinsed with DPBS and then detached by using Accutase (Sigma). The cells were washed three times. Unspecific binding sites were blocked with FACS solution consisting of 2% FBS diluted in PBS for 45 min at room temperature. The cells were then stained with anti CD73-FITC (Invitrogen), anti CD105-APC (Invitrogen) and anti-CD90-PE (Invitrogen) for 1hr at 4°C. The cells were washed 3 times with PBS and kept in FACS solution. Flow cytometry data acquisition was performed in Novocyte2 cytometer and data were analyzed with NovoExpress software.

### Multilineage differentiation of hDPSCs

Dental pulp cells were seeded at a density of 10^5^ cells/cm^2^ and cultured in complete medium until confluency. For osteogenic cultures, medium was composed of A-MEM (Gibco) supplemented with 15% FBS (Sigma), 10^-8^ M dexamethasone (Sigma), 50 µg/ml L-Ascorbate Phosphate (Sigma), 5mM B-Glycerphosphate (Sigma), and 1.8 mM Monopotassium Phosphate (Sigma). Medium was changed twice a week during 3 weeks (28). For adipogenic cultures, complete medium was replaced by A-MEM (Gibco) supplemented with 10 % FBS (Sigma), 10^-5^ M dexamethasone (Sigma), 50 µg/ml L-Ascorbate Phosphate (Sigma), 1 µg/ml Insulin (Sigma) and 0.5 mM isobutylethylxantine (Sigma) (29). Control age-matched cultures were maintained with complete medium in parallel with differentiated cultures for a total of three weeks. Mineralization for osteogenic differentiation was assessed by Alizarin Red staining, whereas adipogenic differentiation was assessed by Oil-Re-O staining.

### Three-dimensional floating sphere culture system

hDPSCs at passage 4 were detached using Trypsin-EDTA (Gibco). About 20,000 cells per well were cultured in ultra low attachment 96 well plate (Thermo Fisher) in neurosphere culture medium (DFNBEb): DMEM/F12(Gibco), 1x P/S (Sigma), 1% N-2 supplement (Gibco), 1% B-27 supplement (Gibco), 20 ng/ml EGF (Gibco), and 20 ng/ml bFGF (Invitrogen) for 7 days. From day 1 to day 3, DFNBEb medium was supplemented with 1 µM SB431542 (Sigma-Aldrich) and 1 µM LDN-193189 (Stemgent). From day 4 to day 7, DFNBEb medium was supplemented with 1 µM SB431542 (Sigma-Aldrich) and 10 ng/ml BMP4 (Stemgent). Half of the medium was replaced every 2 days, and the concentrations were adjusted to final medium volume in culture wells.

### Cell proliferation

Neurospheres formation was monitored daily by taking pictures of selected wells with an inverted microscope (Zeiss, *A*xiovert A1). Diameter measurements were performed by using spheroid sizer tool on MATLAB (30). Data analysis was performed with GraphPad Prism8. Proliferation within neurospheres was assessed by using Click-iT Plus EdU Alexa Fluor 594 kit (Invitrogen) to label the entire population of proliferating cells. Briefly, EdU was first added to the medium at day 0 and renewed at every medium replacement. In some experiments, EdU was added from day 3 and Neurosphseres were collected for immunochemistry analysis following the manufacturer protocol. Population doubling time (PDT) was calculated at day 3 and day 7 *in vitro*. For each experiment, 12 neurospheres were collected in a tube and dissociated with Accutase solution (Sigma) to obtain a cell suspension. The cells were counted manually using KOVACS counting cells. The total number of cells were reported to the number of neurospheres. The PDT was then calculated using the following formula (7):

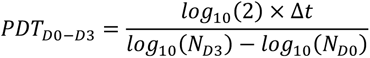

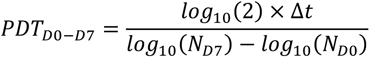

### Otic neurogenic induction of hDPSCs

Otic neuronal differentiation was performed on Gletrex coated surfaces. A 10% Gletrex (ThermoFisher) solution was prepared by diluting the hydrogel matrix in culture medium. Sterile round glass coverslips (ThermoFisher) placed in 4 well culture plate*s* were covered with 180 µl of coating solution. The plates were placed in the incubator at 37°C for 15 to 25 min for Geltrex polymerization.

In order to achieve neuronal differentiation, neurospheres at day 7 *in vitro* were plated on the pre-coated surface. We tested two neuronal induction conditions for a total duration of 14 days *in vitro*. In condition 1, day 7 neurospheres were cultured in DFNBEb medium supplemented with: 0.5 µM ATRA (Sigma-Aldrich), 3 µm CHIR99021 (Stemgent) and 100 ng/ml SHH (R&D Systems) for 5 days, then cultured in a medium supplemented with: 20 ng/ml BDNF (Peprotech) and 30ng/ml NT3 (Peprotech) for an additional 8 days to day 21 *in vitro*. In condition 2, day 7 neurospheres were cultured for 14 days in DFNB medium supplemented with: 20 ng/ml BDNF (Peprotech) and 30 ng/ml NT3 (Peprotech). In both culture conditions, the medium was changed every 2 days. In some culture experiments, the differentiation period was extended for a longer period of maturation up to 32 days in both conditions.

### Co-culture experiments of ONPs and inner ear SGN explants

Cochlear tissue for co-culture was isolated from postnatal day 3 Sprague-Dawley rat pups. 6 rat pups (12 cochleae) were used for each co-culture experiment. Following anesthesia on ice, rat pups were decapitated and the heads rinsed in 70 % ethanol. Under sterile conditions, the skull was opened longitudinally, the temporal bone identified, and the bulla removed and placed into a chilled solution of explant media (comprising 50 mL DMEM (Gibco), 100 µg/ml streptomycin (Sigma-Aldrich) and 100U/ml penicillin (Sigma-Aldrich) and 10mM HEPES). While still submerged in explant media, the spiral ligament and stria vascularis (SV) were removed together by holding with forceps the basal portion of the spiral ganglion (SG) and the SV and slowly unwinding the SV from base-to-apex. We separated the organ of Corti (OC) from the SG and modiolus by holding with forceps the basal portion of the SG and the OC. The SG samples were placed into separate petri-dishes and maintained in chilled explant media until all dissections were completed.

Co-cultures were set up by placing SG explants on Geltrex coated wells for 48 h to ensure adhesion of the explant. The ONPs were then detached using Accutase (Gibco) and co-cultured with SG explants. Cultures were maintained for 5 days in DMEM-F12 medium (Gibco) supplemented with 1% N2 (Gibco), 10 ng/ml BDNF (Peprotech) and 10 ng/ml NT3 (Peprotech) (31). Co-culture samples were maintained in the incubator at 37°C, 5% CO2 and observed daily with Zeiss inverted light microscope. After 5 days of co-culture, the samples were fixed and prepared for immunocytochemistery.

### RNA processing and qRT-PCR

Total RNA samples were collected from all stages investigated *in vitro*. For each stage samples were collected from 3 biological triplicates. The cell samples were lysed with Trizol (Life-science) and RNA purification was done with Zymo kit (R1050, Zymo). RNA quantification was assessed with a Nano Drop 8000 Spectrophotometer (Thermo Scientific). cDNA synthesis was performed using ReadyScript™ cDNA Synthesis Mix (Sigma-Aldrich) in a 20 µl final reaction volume following the manufacturer protocol. For each sample, 400 ng of RNA was used as RNA matrix for the reverse transcriptase enzyme. The cDNA synthesis was performed using 96-well plates in 20 µl final reaction volume using a Bio-Rad C1000 thermocycler according to following program: 5 min at 25°C, 30 min at 42°C and 5 min at 85°C. qPCR was performed using PowerTrack SYBR Green Master Mix (Applied Biosystems, ref. A46012) in 10 µl final reaction volume following the manufacturer protocol.

All qPCR reactions were carried out using 384-well plates in 3 technical replicates in 10 µl final reaction volume on the LightCycler 480 System II (Roche). Reaction mix consisted of Power Track SYBR Green Master Mix 2X, specific primer pairs (Final concentration 0.4-0.8 µM), 1µl of 1:10 diluted cDNA per reaction and H2O to 10 µl. Primer pairs used for gene expression analysis are listed in Supplementary Table 1. The PCR program consisted of an enzyme activation step at 95°C for 2 min followed by 45 cycles of qPCR reaction at 95°C for 5 sec and 60/64°C for 30 sec and finally a melting curve from 60 to 97°C with 5 fluorescence acquisitions per °C.

Expression levels were calculated by the comparative ΔΔCt method (2^−ΔΔCt^ formula), normalizing to the Ct-value of the RPS18 housekeeping gene. For ΔΔCt calculation expressions at day 0 were taken as reference. All values are presented as the mean ± standard deviation. Statistical significance for relative fold change values was determined using one way-ANOVA (*P ≤ 0.05, **P ≤ 0.01, ***P ≤ 0.001, ****P ≤ 0.0001).

### Immunocytochemistry and imaging

Culture samples at different stages of differentiation were fixed with paraformaldehyde 4% for 20 min then washed once with PBS and twice with Tris (Alfaeser) + 0,1% Triton (Sigma-Aldrich). Blocking and permeabilization steps were done with 1% BSA and 0.5 % cold fish gelatin (Sigma-Aldrich) dissolved in Tris + 0.1% Triton solution. Primary antibodies listed in Supplementary Table 2 were diluted in blocking solution and incubated with the samples overnight at room temperature. Then they were washed three times and followed by secondary antibodies incubation for 45 min at RT. DAPI was used to counterstain the cell nuclei. Image acquisition was done on a Leica Thunder microscope. Images were processed with LasX software.

Cells were counted manually using imageJ. Mosaic*s* of imaged wells were used for cell counting. The fraction of immuno-positive cells was reported to the total number of cells stained with DAPI in %. Graphs were established using GraphPad Prism8.

### Nanomechanical characterization by Atomic force microscopy (AFM)

The AFM force spectroscopy experiments were performed using an Asylum MFP-3D head coupled to a Molecular Force Probe 3D controller (Asylum Research, Santa Barbara, CA, USA). Triangular silicon nitride cantilevers (MLCT-NanoAndMore GmbH) with a nominal spring constant of 10 pN/nm were used. The spring constant of the cantilevers was determined using the thermal noise method available within the MFP-3D software. Force-volume maps were performed in PBS buffer, at room temperature and with a maximum loading force of 1 nN, recording at least 10 maps per cell type. The recorded AFM-FS data were used to determine the Young’s modulus (E) of the tissue using a modified Hertz contact model (1, 2). Histograms of the distribution of Young’s moduli were fitted with a Gaussian function to obtain the mean E value of the probed cells. One-way ANOVA was used for comparison of Young’s modulus values. Tukey’s multiple comparisons were employed after performing normality tests for comparison of every mean with every other mean. Three cell types were studied (hDPSCs, native SGN and hDPSC-derived SGN-like cells) under live and fixed conditions. Statistical significance was set at p ≤ 0.05.

## Results

### Cells isolated from wisdom teeth pulp tissue have mesenchymal stem cell characteristics

The isolated hDPSCs have an elongated shape at the onset of culture. These cells have the ability to form colonies, with high adherence and proliferation activities. The subcultured cells exhibited flattened and fibroblastic morphology and are virtually all STRO-1 immuno-positive (**Figure 1A**). We also demonstrated that the isolated hDPSCs were immuno-positive for the known mesenchymal antigens CD73, CD90 and CD105 using flow cytometry. The flow cytometry analysis showed that more than 96% of the cells co-expressed CD90, CD73 and CD105 markers (**Figure 1B**). Furthermore, we confirmed the previously reported multilineage capacity of hDPSCs to be converted to osteogenic (29) and adipogenic (32) cell lineages. Their potential to differentiate into osteoblasts was confirmed by the production of calcium deposits via Alizarin Red staining after 3 weeks of induction (**Figure 1C**). Adipogenic differentiation was demonstrated by Oil-Red-O staining of lipid droplet after 3 weeks of induction *in vitro* (**Figure 1D**). The cultured hDPSCs maintained in osteogenic medium showed a significant absorbance increase (3-fold) (**Figure 1E**), while those maintained in adipogenic medium exhibited about 2-fold increase absorbance (**Figure 1F**) when compared to untreated age matched-control cells.

**Figure 1.**
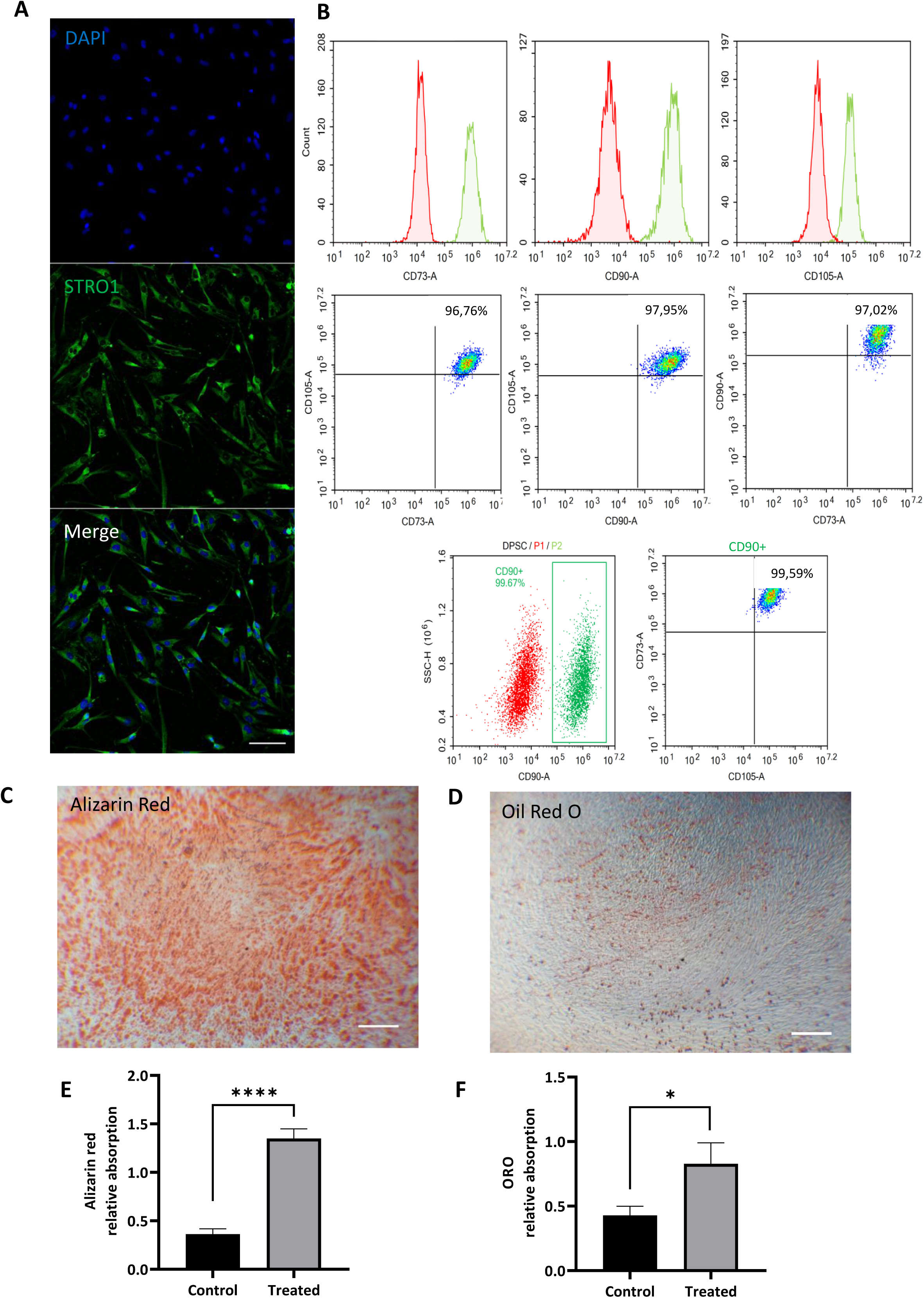
Identification of mesenchymal stem cell markers and multilineage charaterization of the isolated hDPSCs. **A.** hDPSCs from passage 4 were immunostained with an antibody against Stro1 (green). Stro1 immunostaining was detected in virtually all the undifferentiated hDPSCs in culture. Nuclei were stained with DAPI (shown in blue). Scale bar = 100 µm. **B.** Analysis of undifferetiated hDPSCs by flow cytometry indicates the ratio of immunopositive cells for the known mesenchymal characteristic markers: CD73, CD90 and CD105. More than 96% of hDPSCs express either CD73 or CD90 or CD105. Approximatively 99% of CD90+ cells express are CD73 and CD105 positive.

### Neurosphere formation

It was previously demonstrated that culturing hDPSCs as 3D-aggregates could drive them into relatively less heterogenous cell subpopulations (33–35). Cells initially expanded as a monolayer were passaged and seeded under floating culture condition (**Figure 2A)** and aggregated into neurospheres as outlined in **Figure 2B**.

**Figure 2.**
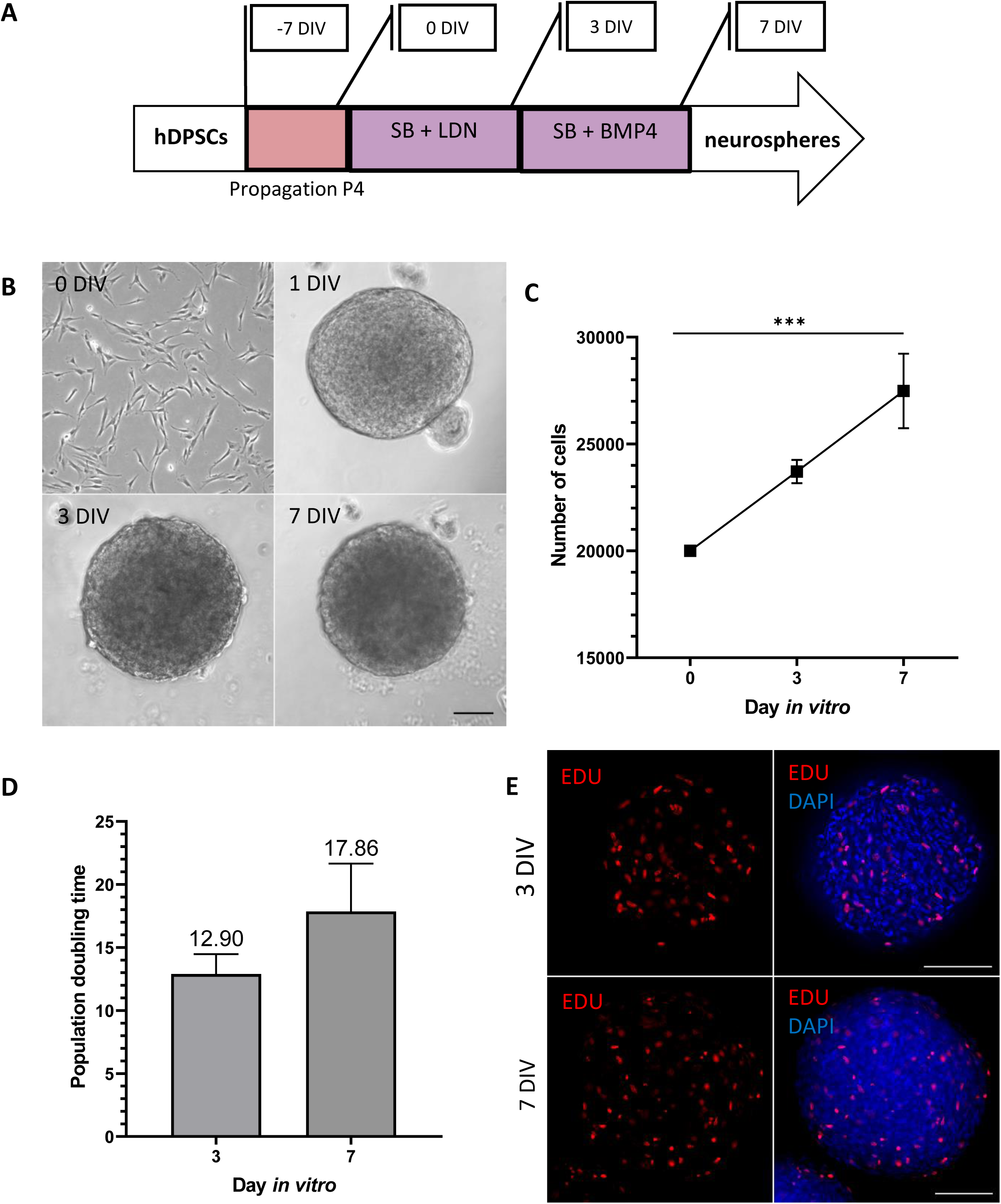
Morphometric analysis of hDPSC-derived neurospheres. **A.** Schematic illustration outlining the timeline and conditions of the newly established protocol for the generation of hDPSC-derived neurospheres. The cells were exposed to SB/LN for 4 days *in vitro* followed by treatment by SB/BMP4 for additional 3 days *in vitro*. Abbreviations, SB: SB431542: TGFb inhibitor, LDN: inhibitor of the bone morphogenetic protein (BMP) pathway inhibitor, BMP4: Bone mophogneic protein 4. **B.** Phase contrast images demonstrate the ability of dissociated hDPSCs to aggregate in floating three-dimensional spheres, as early as 24 h after cell seeding. The generated 3D spheres have semi-transparent appearance. At day 0, cells are in a homogeneous suspension. After 24 h, a 3D sphere is formed by cell aggregation in a medium supplemented with N2 +B27+ bFGF+ EGF. **C.** Quantification of the number of cells per neurosphere. Cells counting demonstrated a progressive increase in cell number at different time points (**day 1 to day 7**) during neurosphere generation to reach ∼27.5×10^3^ ± 3000 cells at day 7 *in vitro*. **D.** Evaluation of proliferation ability within the neurospheres by population doubling time (PDT). The cells demonstrated a steadily rising growth with a more evident increase from day 0 up to day 3 *in vitro and* resulting in a PDT of 12.90 (±3.15) and then to a PDT of ∼ 17.86 (±7.5) at day 7 *in vitro*. **E.** Assessment of proliferation using Click-it EDU supplied in the culture medium during the period of sphere formation. The EdU is incorporated in the floating spheres with high proliferative capacity. Edu was supplemented in culture medium between day 0 and day 3 then between day 3 and day 7 *in vitro*. Staining indicates that proliferation occurs in both phases of culture. Statistical test T-test was used **P≤ 0.001, n=4. Scale bar= 100 μm.

An equal number of cells from each initial culture condition were transferred to low attachment culture plates, and typical characteristics of neurosphere formation (i.e., number of cells, PDT, viability, and proliferation) were assessed after 3 and 7 days in cell culture (**Figures 2C-E & S1**). The immature cells at day 0 *in vitro* were taken as controls. Overall, in a typical culture composed of approximately 2 x 10^4^ seeded cells/well, the dissociated cells aggregate, and the neurospheres are formed after 1 day of culture (**Figure 2B**).

Proliferative activity was observed within the neurospheres at different stages of the culture (**Figure 2C-E**). Proliferation was confirmed by the increased number of cells/neurosphere that reached 27.481 cells ±3400 per neurosphere at day 7 *in vitro* (**Figure 2C)**. Proliferation was characterized by an increased PDT during the time course of differentiation from ∼12.90 (±3.15) at day 3 to ∼17.86 (±7.5) at day 7 *in vitro* (**Figure 2D**) and Edu staining (**Figure 2E**).

### The two-step neurosphere assay is more efficient for expression of neural crest and otic placodal markers

To specifically assess the fate of differentiated cells during the early phase of differentiation, cells were either collected or fixed at day 3 and day 7 *in vitro* and analyzed by qPCR and immunocytochemistry, respectively, for specific cell type markers of neural crest (NC) and otic lineage specification i.e., otic placode (OP), otic vesicle (OV) and otic neuronal progenitors (ONP) (**Figure 3A-B**).

**Figure 3.**
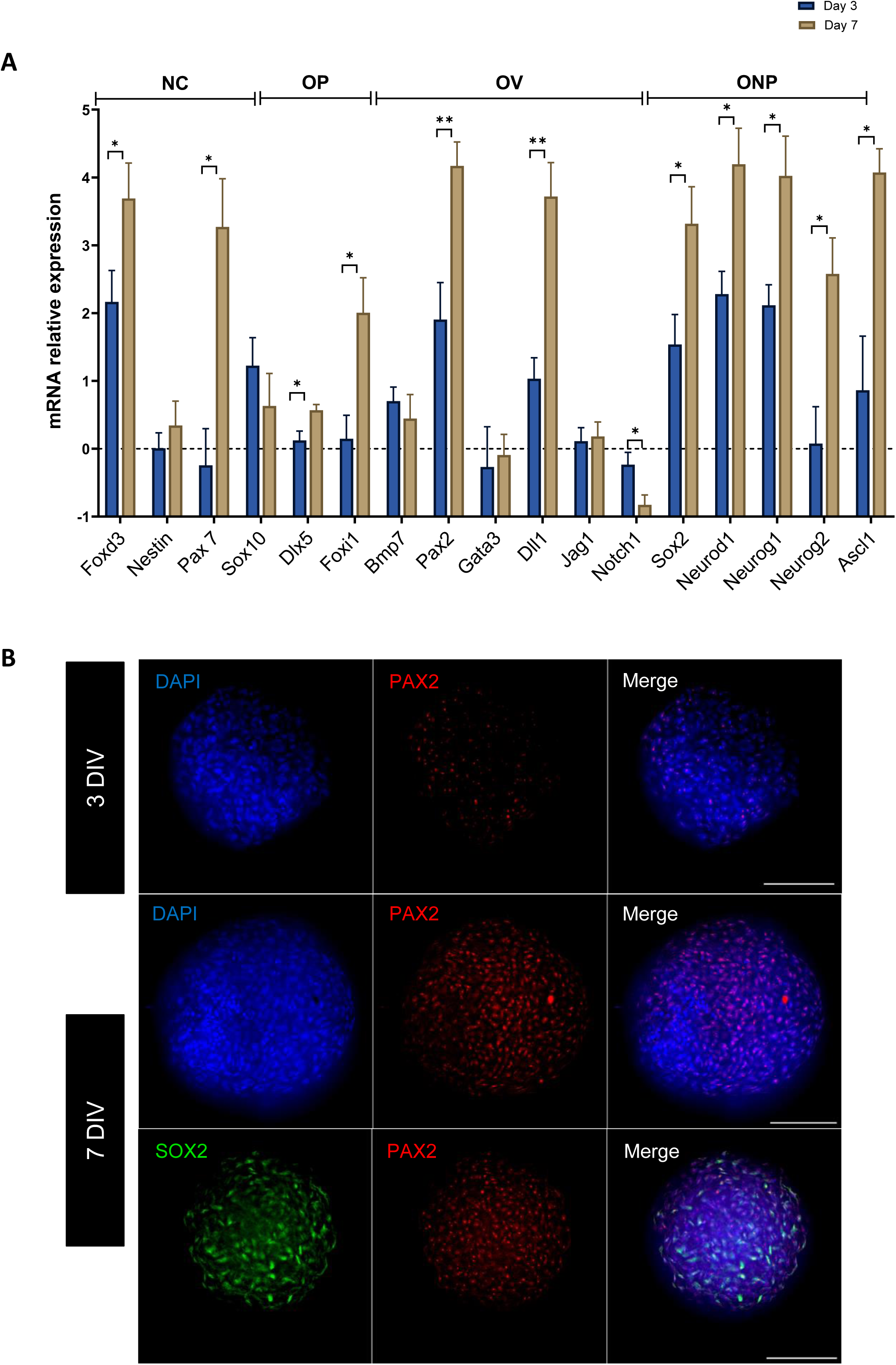
Induction of expression of early neuronal and otic/placodal markers in hDPSC-derived neurospheres. **A.** Bar chart showing the relative gene expression levels in logarithmic (Ln) scale obtained by qPCR analyses for four distinct panels of genes featuring the neural crest, otic placode/vesicle, and otic neuronal progenitors, respectively. Cells were collected at day 3 and day 7 of differentiation of the protocol (A) and analyzed to assess the impact of the dual inhibition/activation of BMP signaling under continuous TGFb inhibition on otic induction. Noticeably, results demonstrate a significant upregulation of otic/vesicle and otic neuronal progenitor markers (ie., *Pax2, Sox2, NeuroD1, Neurog1*) in the SB/LDN/BMP4 culture condition, which is the condition that yielded optimal otic induction from hDPSCs. Statistically significant differences as compared to undifferentiated cells at day 0 are indicated by *P≤ 0.05, **P≤ 0.01 (n=3 one way ANOVA), n=3 independent experiments. **B.** Representative images showing PAX2 and SOX2 immuno+ cells at day 3 and day 7 during the time course of differentiation. PAX2 was detected from Day 3 however SOX2 expression were detected only from day7. Cell nuclei were counterstained with DAPI (blue). Scale bars = 100 µm.

We explored the dynamic expression of a comprehensive panel of preselected OP/OV and ONP known gene markers. Noteworthy, qPCR analyses showed that treatment with SB/LDN for 3 days and then SB/BMP4 for 3 days led to a robust expression of NC progenitor cell markers (Sox10, FoxD3) and OV/OP markers, such as Pax2 (**Figure 3A**). During inner ear development, OP induction is swiftly followed by OV formation. OP marker expression analysis at 7 days of differentiation demonstrated a significant upregulation of a subset of genes such as Pax2/Dll1, which are among key otic/placodal markers. Importantly, significant upregulation of ONP-associated genes (i.e. Neurog1, Neurod1, Ascl1) was detected at day 7 when compared to immature cells at day 0 and early differentiated cells at day 3 *in vitro* (**Figure 3A**). Moreover, neural crest markers such as FoxD3 and Pax7 were also significantly higher at day 7 when compared to day 3 cell cultures.

To support the qPCR results, we also studied in parallel the expression of neural crest (NESTIN) and ONP (SOX2 and PAX2) markers by immunohistochemistry analysis. NESTIN immunoreactivity was similar between day 3 and day 7 neurospheres. About 95% of cells remained immunopositive for the expression of NESTIN in neurosphere-derived hDPSC sub-populations (**Figure S2**). The expression of PAX2 and SOX2 at day 3 as compared to day 7 *in vitro* indicated an early expression of PAX2 from day 3, whereas SOX2 expression was only detected by day 7 *in vitro* (**Figure 3B**). Both SOX2 and PAX2 are colocalized at the nuclear level (**Figure S3).** These examples of immunolocalization of neural crest and ONP markers at days 3 and 7 in differentiated neurospheres support overall data from qPCR analysis. Altogether, these data from qPCR and immunohistochemistry analyses demonstrate that the differentiated cells from hDPSCs have rapidly engaged toward OP/OV and ONP fate by day 7 *in vitro* upon SB/LDN and SB/BMP4 treatment using a neurosphere culture system.

### Enrichment of DPSC-derived cells expressing ONP and to SGN early markers

Previous studies have demonstrated a role of Shh, RA and WNT signaling pathways in otic neuronal development *in vivo* (36–39). Additionally, two crucial neurotrophins (i.e., BDNF and NT-3) and their associated receptors, TrkB and TrkC have been shown to be necessary for the normal development of afferent innervation during early human inner ear development (21,40,41). All of these signaling pathways and neurotrophin molecules can also enhance the production of SGN-like cells from pluripotent stem cells in mammalian 3D inner ear organoids (23,24,42,43). Thus, we tested whether prolonging the culture period under exposition to WNT/SHH/CHIR for 5 days followed by BDNF/NT3 for 8 days *in vitro* (paradigm1) could promote the cells into more differentiated otic neuronal lineage compared to treatment with BDNF/NT3 alone for the equivalent period (paradigm 2) (**Figure 4A**). The DPSC-derived neurosphere treated cultures underwent rapid and profound morphological changes leading to neurite outgrowth from differentiating cells at the border of the neurosphere, suggesting their progressive neuronal differentiation (**Figures 4B-C**). Of interest, a subset of DPSCs exposed to either neuronal inductive paradigm 1 or 2 acquired a bipolar morphology consistent with potential early differentiation of otic neuronal progenitors (ONPs) and SGN-like cells during the period from day 14 to day 21 *in vitro*.

**Figure 4.**
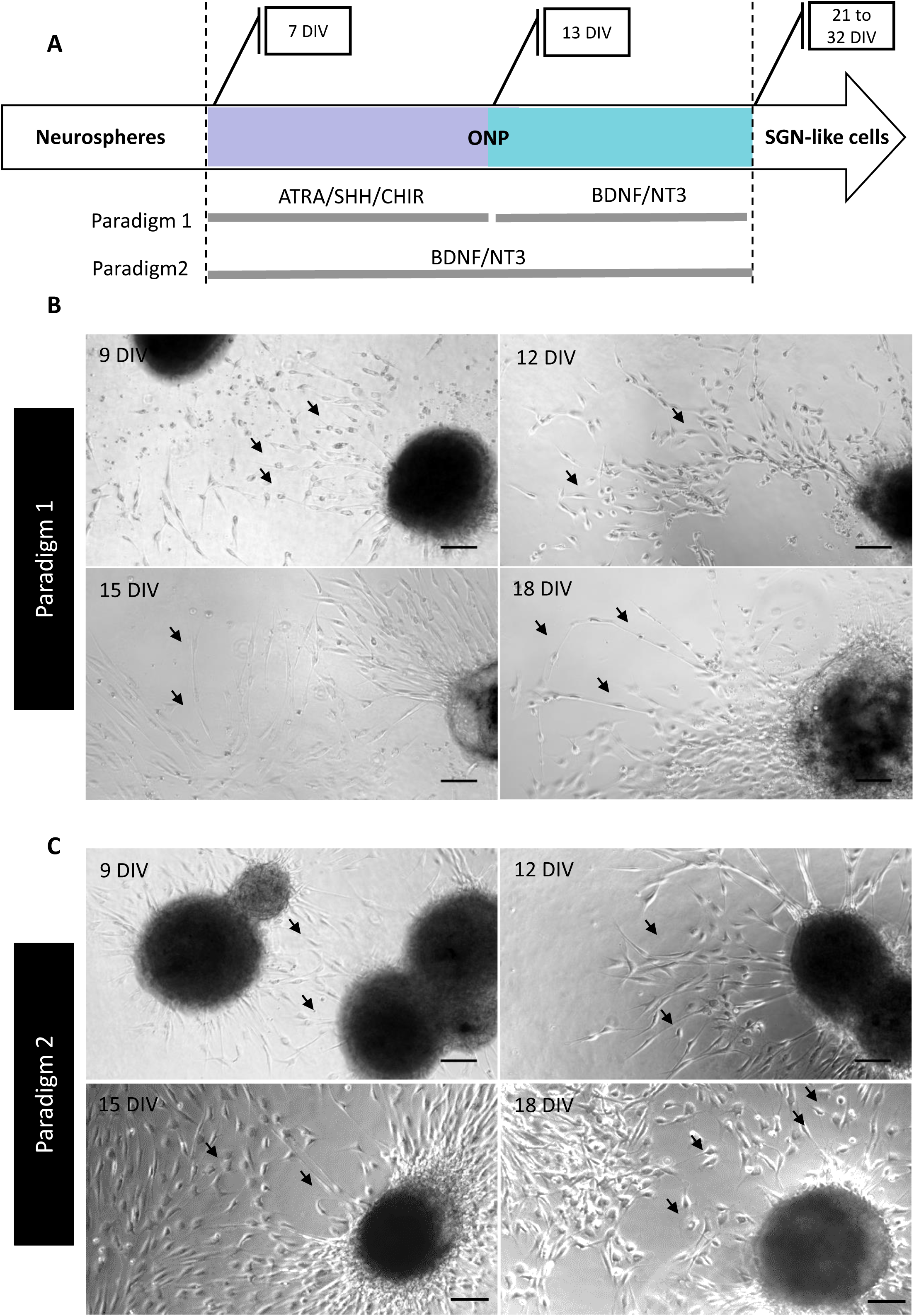
Differentiation culture conditions post neurosphere generation. **A**. Timeline of differentiation and experimental paradigms. Neurospheres were derived from hDPSCs before being allocated into paradigms 1 or 2 for a further 15 days *in vitro*. **B-C**. Phase-contrast representative images showing the morphological characteristics of hDPSC-derived otic neuronal progenitors during the time course after neurosphere generation. In **B** the differentiated cells displayed homogeneous otic neuronal progenitors with round soma migrating outside the neurosphere borders after treatment with ATRA/SHH/CHIR factors, then elongating. In (**C**) the differentiated cells displayed heterogeneous elongated otic neuronal progenitors after treatment with BDNF/NT3 neurotrophins. Scale bars = 100 µm.

Under culture paradigm 1, we observed a noticeable homogeneity of cells migrating from the neurospheres (**Figure 4B**). These cells are characterized by a round shaped soma between day 9 and day 12 *in vitro,* then a progressive elongation toward a bipolar morphology. However, under culture paradigm 2, we noticed a less homogeneous cell population (**Figure 4C**). These cells form a dense network of elongated cells around the neurospheres. Moreover, the plated neurospheres maintained a preserved structure in paradigm 2 when compared to that displayed in paradigm 1.

Investigating the variations in the relative expression levels of some key gene markers allowed for a comparison between neuronal induction paradigm 1 and 2 at day *14 in vitro*, revealing that the main difference is related to the expression levels of a subset of ONP markers **(Figure 5A)**. At day 14 *in vitro,* paradigm 1 induces a higher expression of Bmp7 when compared to its level in paradigm 2 culture condition. The expression of Bmp7 in combination with Pax2 and Sox2 supports that paradigm 1 enhances the ONP phenotype. In the same way, paradigm 2 also induces the expression of ONP related genes i.e., Bmp7, Pax2 and Sox2. However, paradigm 2 induces a significantly higher expression of Neurod1 and Trkb with Prph, suggesting strong early commitment toward SGN-like cells phenotype under this culture condition. This suggestion could be supported by the significant downregulation of the expression of dlx5, which is related to the phenotype of early otic/placode cell progenitors. Moreover, the expression of PAX2 and PRPH **(Figure 5B)** in both culture paradigms confirms the ONP lineage induced by both paradigms 1 and 2 at 13 days *in vitro*.

**Figure 5.**
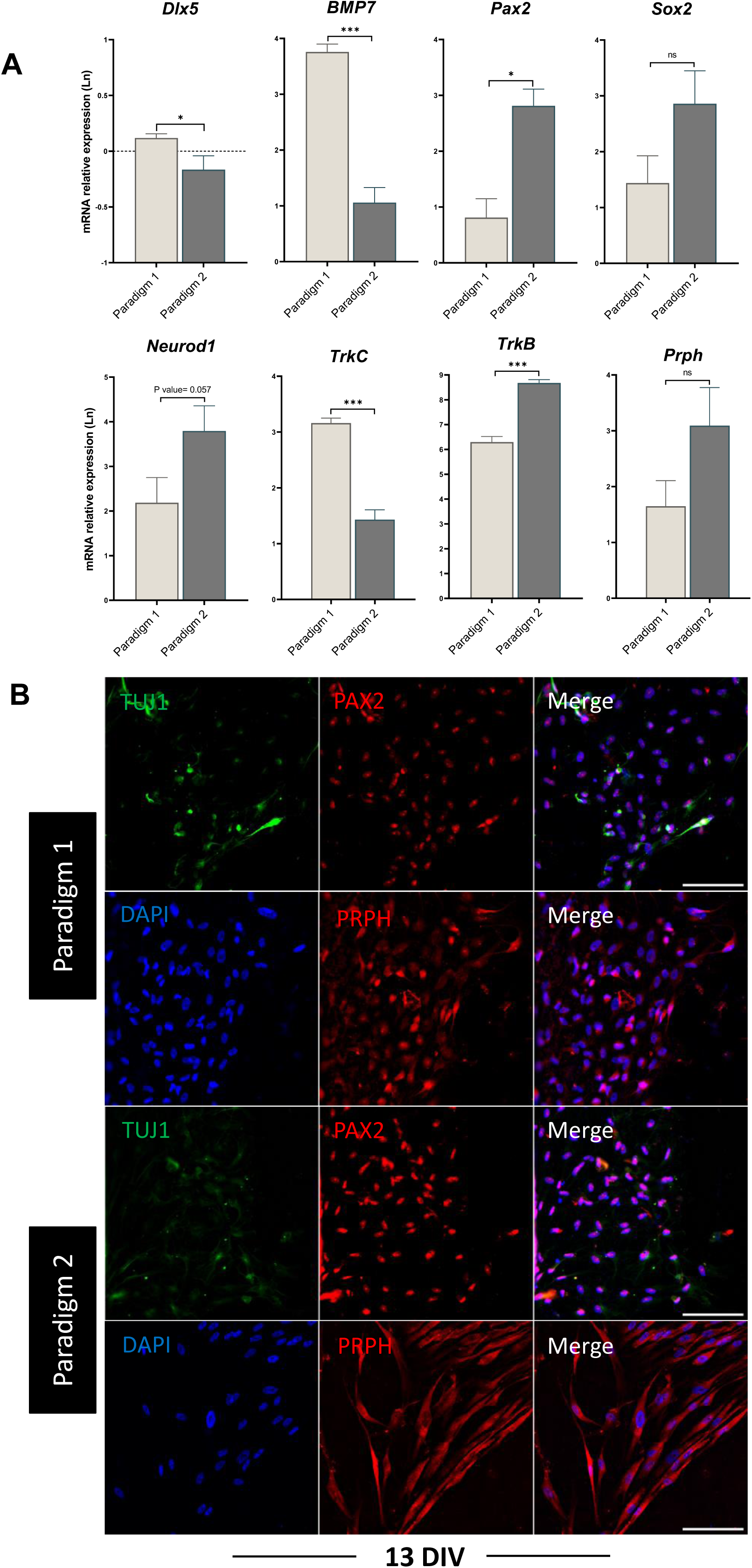
Characterization of ONP phenotype in differentiated cells at 13 DIV. By using this protocol, neurospheres were exposed to either ATRA/SHH/CHIR until day 13 *in vitro* and to BDNF/NT3 until day 21 i*n vitro (paradigm 1) or only exposed to* BDNF/NT3 for the same period *in vitro (paradigm 2)*. **A-** Bar charts showing the relative gene expression levels in logarithmic (Ln) scale obtained by qPCR analyses for a panel of ONP related lineage from paradigm 1 (Figure 5B) and from paradigm 2 (Figure 5C). Cells were collected at 13 DIV. Results indicate the induction of several genes related to ONP phenotype. **B-** Expression of TUJ1 (green), PAX2 (red), PRPH (red) in ONP from paradigms 1 &2 at 13 DIV. Statistical differences were determined with T-test. P values are indicated with *P≤ 0.05, n= 3. Scale bar = 100 µm.

At day 21 *in vitro*, qPCR analysis of differentiated cells from both culture paradigms (**Figures 6A-B**) showed strong evidence of the ongoing otic differentiation process. The expression of a panel of NC and OP gene markers was significantly downregulated in both culture conditions (i.e., Nestin, Snail, Eya1, Six1). The expression of SGN-related genes (Sox2, Neurod1, Neuorg1) in both paradigms showed a slight decrease but was not statistically significant. Of interest, the expression of TrkB, a marker of neuronal maturation, increased significantly under both *in vitro* differentiation paradigms.

**Figure 6.**
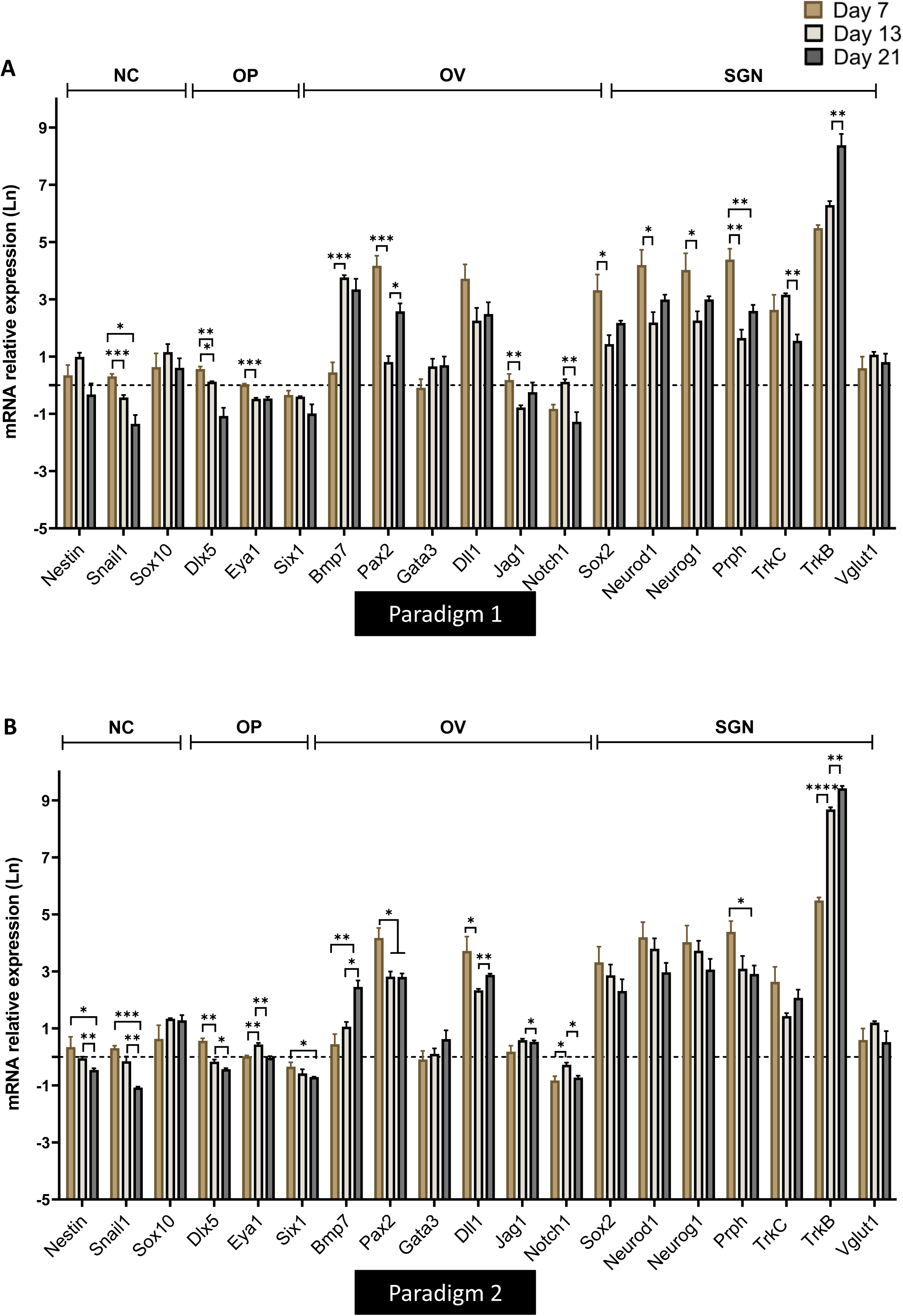
Analyses of otic placode/vesicle and SGN gene markers in differentiated cells using our newly established protocol outlined in. **figure 5A**. By using this protocol, neurospheres were exposed to either ATRA/SHH/CHIR until day 13 *in vitro* and to BDNF/NT3 until day 21 i*n vitro (paradigm 1) or only exposed to* BDNF/NT3 for the same period *in vitro (paradigm 2)*. **A-B**. Bar charts showing the relative gene expression levels in logarithmic (Ln) scale obtained by qPCR analyses for distinct panels of genes featuring NC, OP/OV and SGN lineages from paradigm 1 (Figure 5A) and from paradigm 2 (Figure 5B). Cells were collected at day 7 of differentiation, day 13 and day 21 of differentiation according to paradigms 1 and 2 of the procedure. For each time point, the samples are collected 3 independent experiments. Results indicate a downregulation in the expression of NC and OP markers whereas SGN related markers remained expressed ie Neurod1, Prph, TrkC. TrkB is the marker that showed the most significant upregulation in both conditions. Statistical differences were determined with one way ANOVA. P values are indicated with *P≤ 0.05, **P≤ 0.01, ***P≤ 0.001, ****P≤ 0.0001.

We next examined the expression of a subset of mature neuronal lineage (i.e., TUJ1, MAP2, NEUN, SOX2) and SGN related (PRPH, TRKC), markers (data not shown) on differentiated cells from both paradigms at day 21 *in vitro* by immunohistochemistry (**Figure 7 A-B**). In both culture conditions, differentiated cells expressed neuronal markers such as NEUN, TUJ1 and MAP2. These differentiated cells have a bipolar morphology close to that of native SGNs *in vivo*. To gain further insights into the differentiation state of SGN-like cells at day 21 *in vitro*, we quantified the number of SOX2, NEUN and GFAP immunopositive cells in both culture paradigms (**Figures 7C**). We found that the number of SOX2 immunopositive cells was significantly higher in culture paradigm 1 (about 80% of total cells) when compared to culture paradigm 2. In contrast, the number of GFAP immunopositive cells was higher in culture paradigm 2 (about 40.7% of total cells).

**Figure 7.**
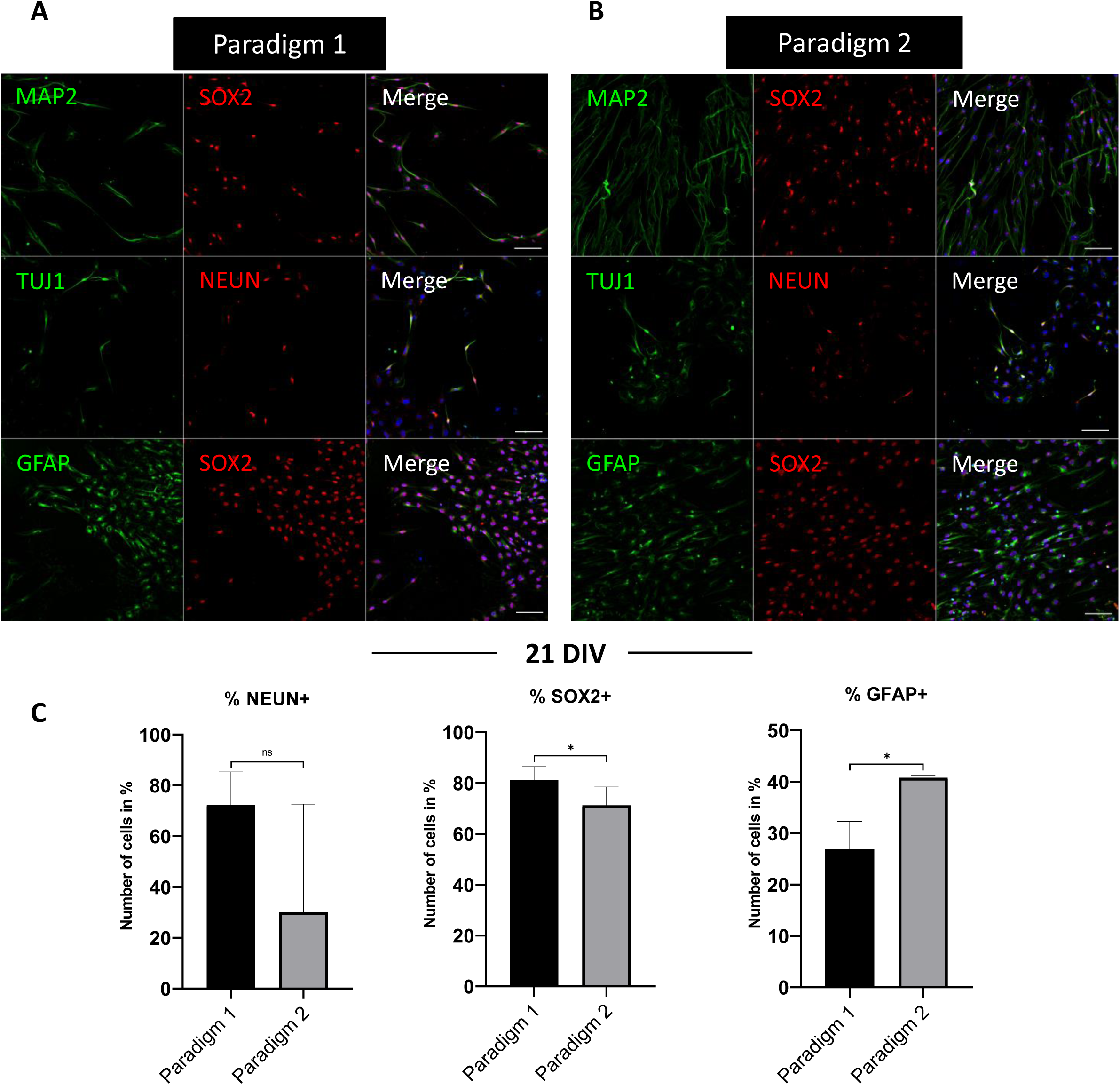
Representative images of immunocytochemical analyses and quantification of the expression of neural markers in cultures at 21 DIV in paradigms 1 and 2. **A.** Expression of MAP2/TUJ1/GFAP (shown in green) and SOX2/NEUN (shown in red) in paradigm 1 cultures. **B.** Expression of MAP2/TUJ1/GFAP (shown in green) and SOX2/NEUN/ (shown in red) in Paradigm 2 cultures. DAPI staining is shown in blue. Scale bars, 100 μm in all panels. **C.** Quantitative analysis revealed a greater number of SOX2 (80 % of total) in paradigm 1 and NEUN (40% of total) immuno+ cells in paradigm 2 at 21 DIV. Statistical differences were determined with T-test. P values are indicated with *P≤ 0.05, n=3. Scale car = 100µm.

When the differentiation period was extended to 32 days *in vitro* (DIV) (**Figure 8A**), we observed that only the cultures differentiated under paradigm 2 were maintained with a fairly good cell viability as compared to cells differentiated in paradigm 1. The advanced cultures at 32 DIV of paradigm 2 exhibited a more mature bipolar phenotype with multiple neuronal populations. Interestingly, about 40 % of the bipolar SGN-like cells were BRN3A/SOX2 immunopositive (**Figure 8B),** in addition to TUJ1, PRPH and TRKC. The expression of these key otic neuronal lineage markers in differentiated cells at this stage of culture (i.e., 32 DIV) may suggest their potential differentiation toward SGN-like cells.

**Figure 8.**
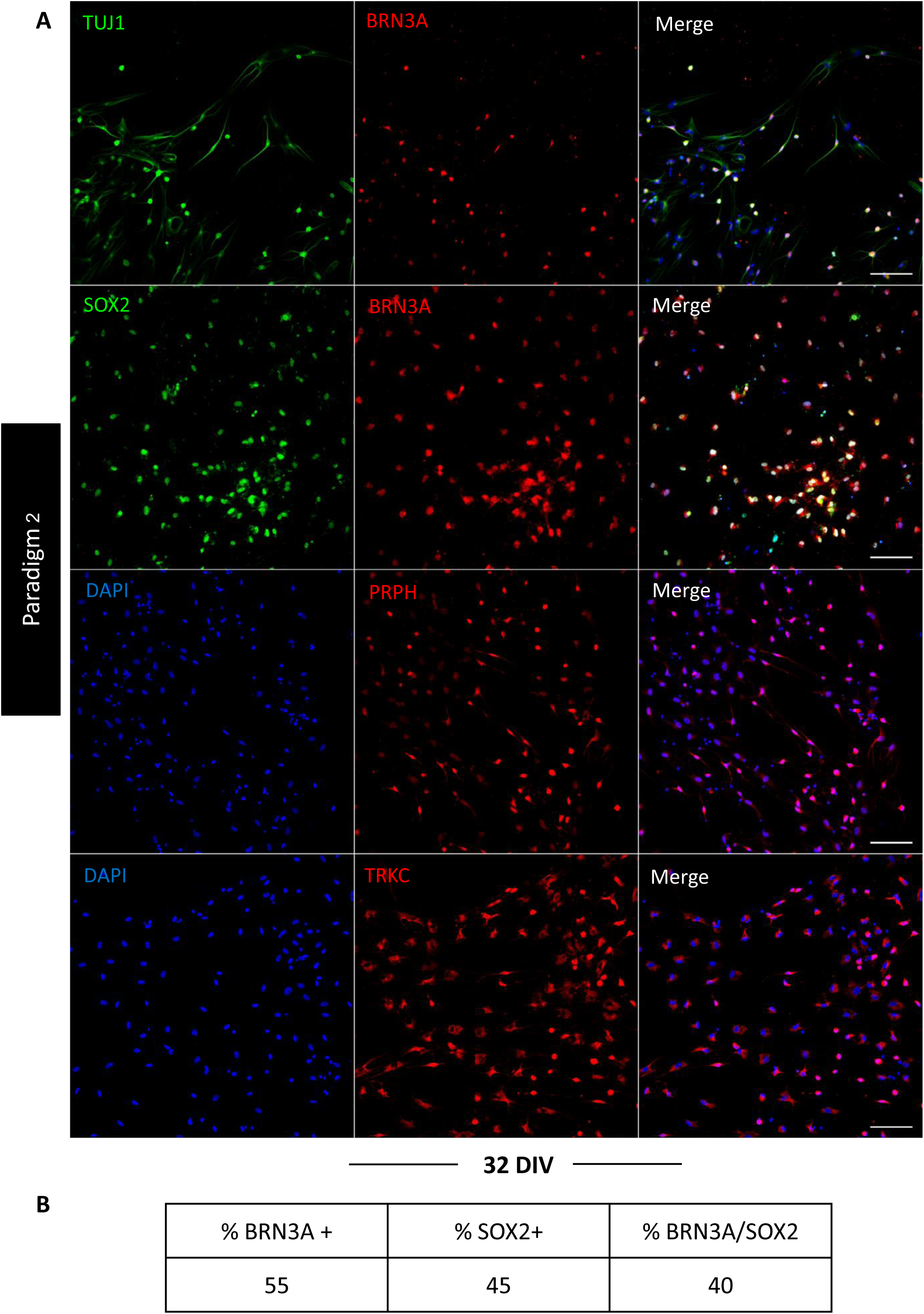
Characterization of the expression of SGN markers at 32 Div in paradigm 2. **A.** Among the differentiated neuronal cells SGN cells are bipolar and express SOX2, BRN3A, TUJ1, PRPH and TRKC.. Scale bar= 100 µm **B.** The table represents the purcentage of cells expressing SOX2, BRN3A and SOX2/BNR3A. A total of 2700 cells was counted.

### AFM characterization revealed similarities between SGN in vitro and SGN in vivo

To add a new dimension of characterization, we explored the nanomechanical properties of our generated SGN like-cells derived from culture paradigm 2 at 32 DIV and compared them to *in-vivo* SGN from rat pups and undifferentiated hDPSCs. The results showed that both SGN like-cells and *in vivo* SGN share the same Young’s modulus in either fixed samples (5-10 KPa) (**Figure 9A**) or unfixed samples (∼0.5-0.7 KPa) (**Figure 9B**). These measurements of Young’s modulus were significantly lower than the measurement from undifferentiated hDPSCs which were stiffer. Interestingly, topographic reconstructions revealed that both *in vivo* SGN and *in vitro* SGN share a bipolar elongated morphology (**Figure 9 C-D**) where hDPSCs are characterized by their known elongated fibroblastic morphology (**Figure 9E**). The evident contrast between *in vitro* SGN-like cells and the hDPSCs from where they were derived, in addition to the major difference between their nanomechanical properties, are strong sign of the otic neuronal differentiation process.

**Figure 9.**
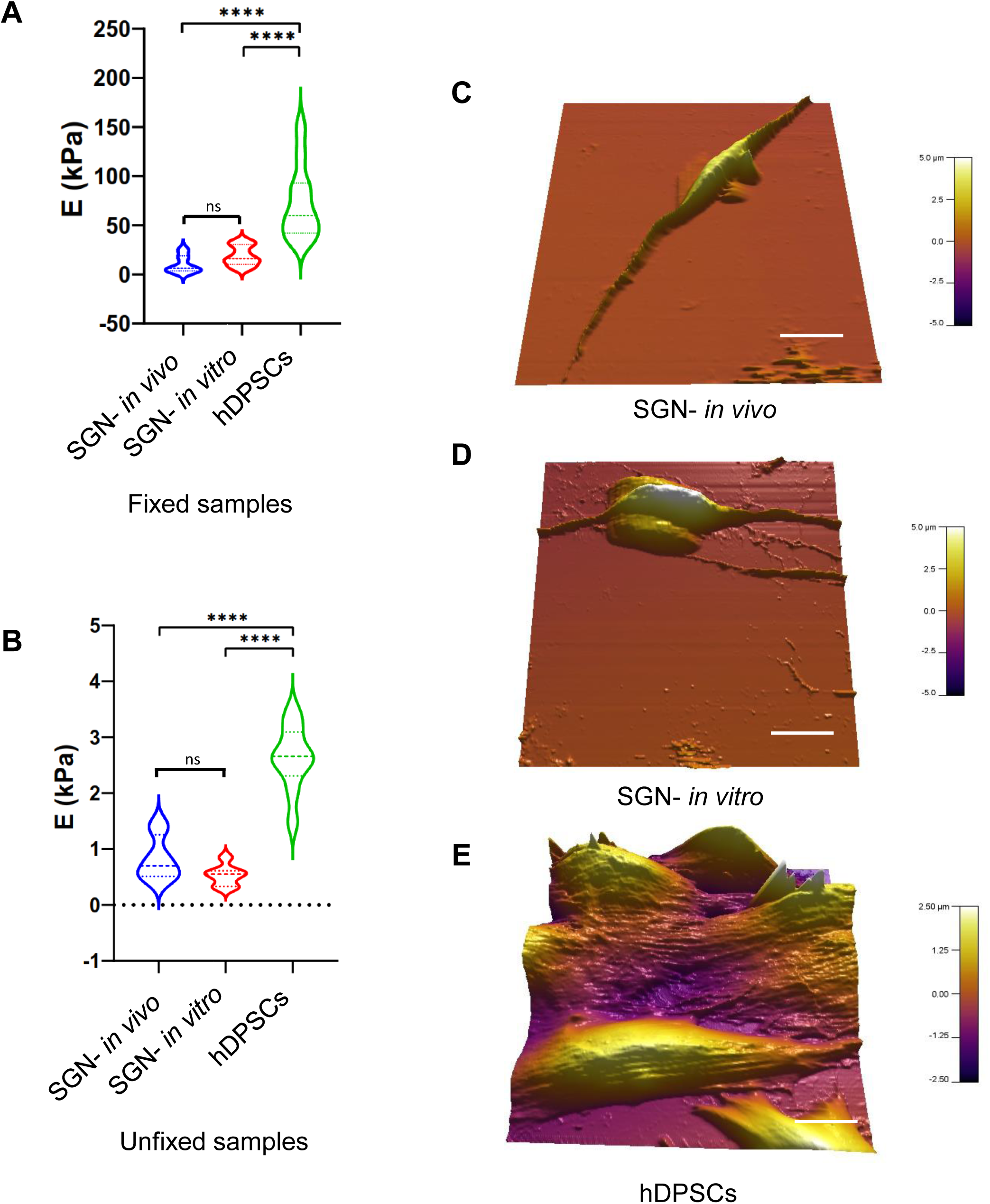
Nanomechanical characterization of SGN-like cells at day 32 *in vitro* by AFM. **A.** Violin Plot of measured Young’s modulus of fixed SGN *in vivo*, SGN *in vitro*, and hDPSCs. **B.** Violin Plot of measured Young’s modulus of unfixed SGN in vitro, SGN in vivo and hDPSCs. **C.** 3D reconstruction of analyzed SGNs *in vivo*. **D.** 3D reconstruction of analyzed SGN *in vitr*o. **E.** 3D reconstruction of analyzed hDPSCs. SGN *in vivo* are like SGN *in vitro* and different from hDPSCs in terms of shape or nanomechanical properties. Statistical differences were determined with one way ANOVA. P values are indicated with ****P≤ 0.0001. n= 10 measurements. Scale bar = 20 µm.

### ONP promotes SGN neurite outgrowth in co-culture

To investigate the therapeutic potential of our ONPs in restoring SGNs, we co-cultured ONPs from paradigms 1 & 2 at 13 DIV with SGN explants from rat pups. We observed neuronal growths of SGNs **(Figure 10A)** which was preferentially directed toward the ONPs (**Figure 10B**) and, in some co-cultures, this neurite extension was beyond 1000 µm. Moreover, we noticed that these neurites made contacts with the ONPs at the level of their membranes. To confirm these new contacts between ONPs and neurites projecting from the SGN explant, we performed a 3D reconstruction using Imaris software. Interestingly, we were able to clearly observe the neurites that were in direct contact with the membranes of the ONPs (**Figure 11**), which is strong evidence of the potential of ONPs to reconnect with the SGN extracted from the inner ear of the postnatal rat. This observation led us to hypothesize that ONPs could potentially secrete trophic factors and chemokines that may affect neurites outgrowth, considering the rich composition of hDPSCs secretum with neurotrophic factors and growth factors, such as NT3 and IGF.

**Figure 10.**
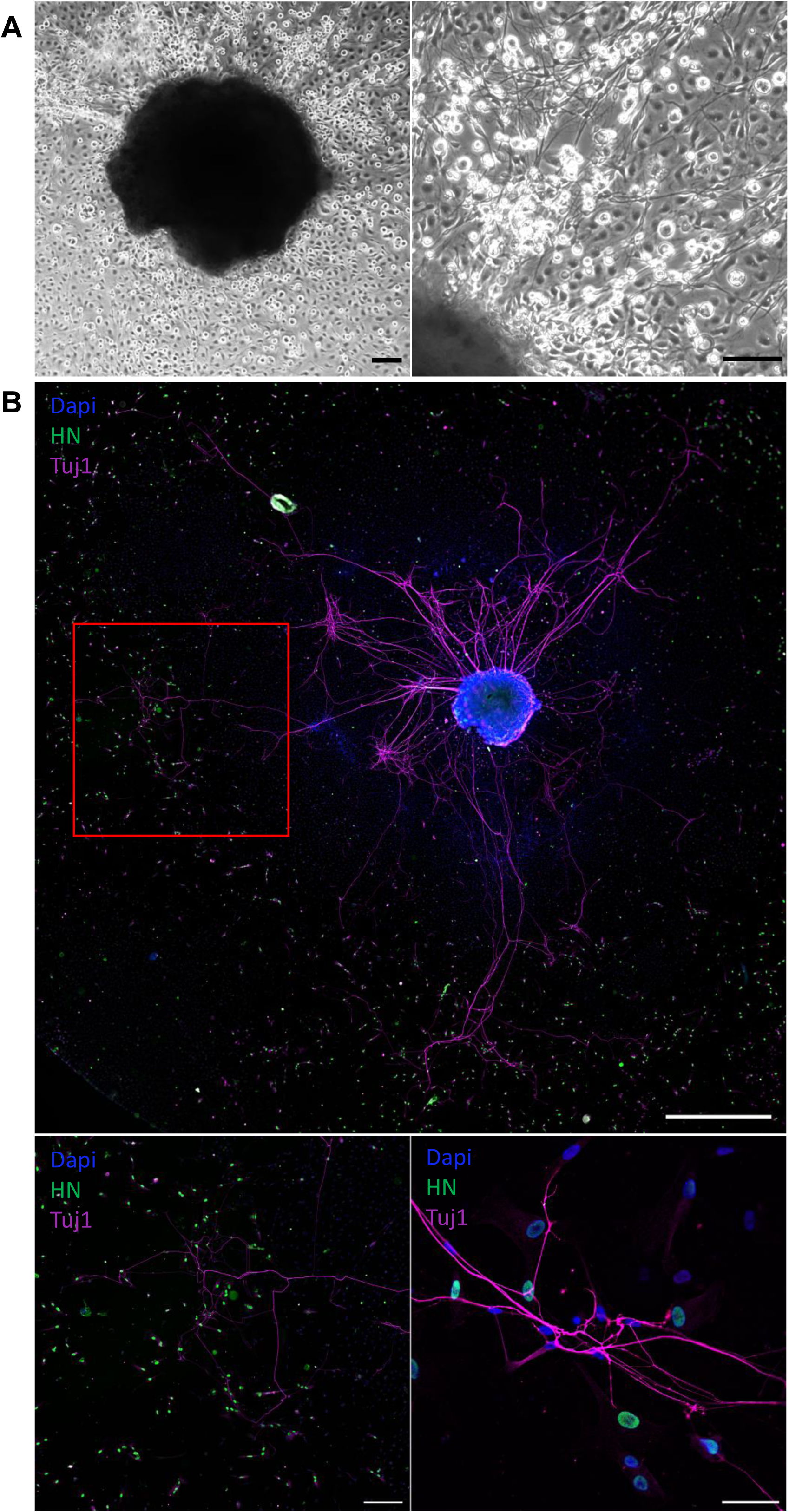
Characterization of the co-cultures between ONP and SGN explant. **A.** Representative image of the co-cultured explant of SGN under phase contrast microscope at 7 DIV. The images show neurons that migrated from the explant with neurite outgrowth toward the ONP. **B.** Representative images of neurite outgrowth (stained with Magenta= TUJ1) that project toward the ONPs (green= human nuclei & Magenta = TUJ1). DAPI was used to counterstain the nuclei.

**Figure 11.**
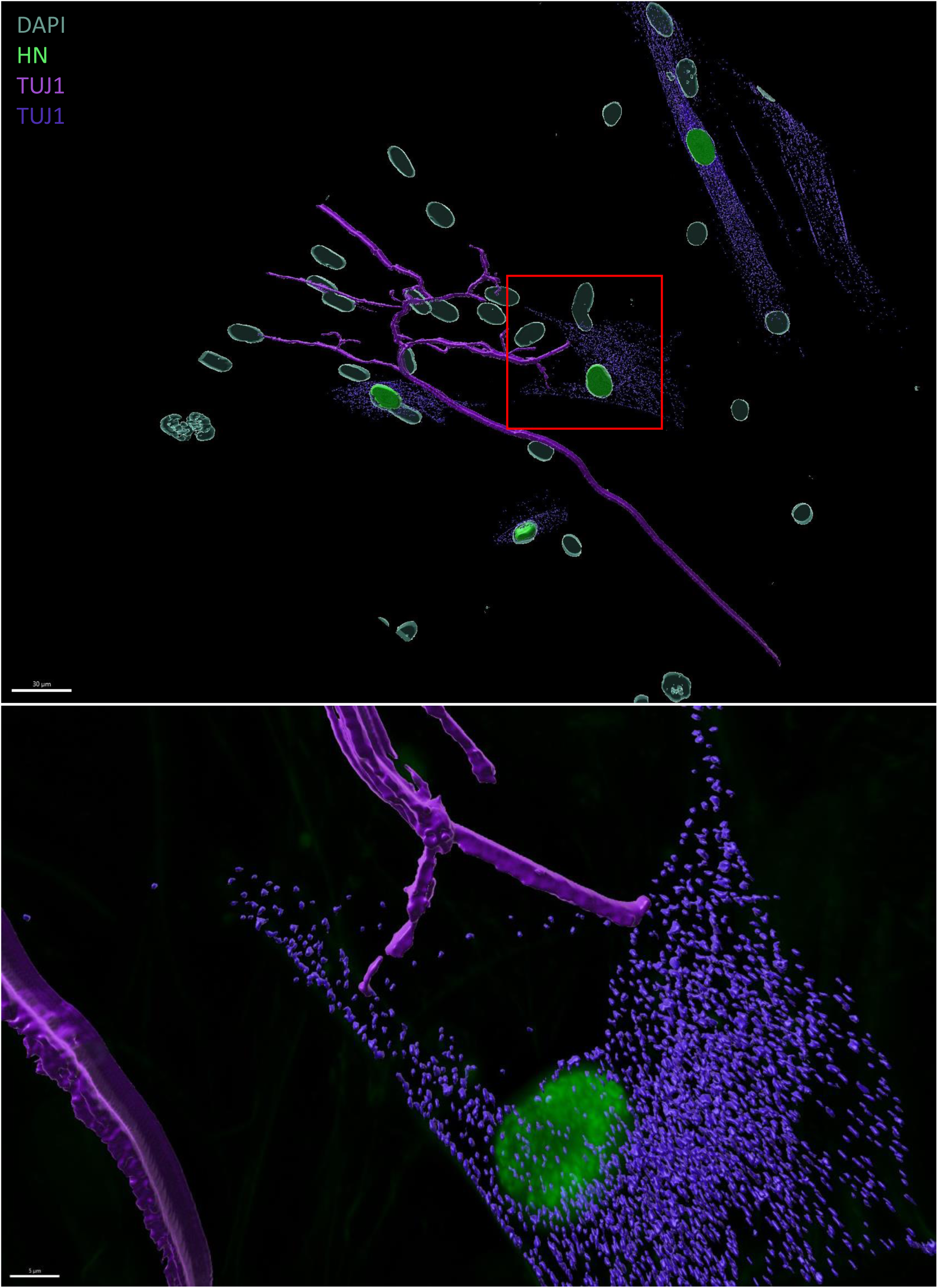
3D reconstruction of neuritic contacts with ONP using IMARIS. Images represent the 3D reconstruction of neurites contacts with the ONPs using IMARIS software. The neurites (purple= TUJ1) make contacts with the membrane of the ONPs (green= human nuclei & purple = TUJ1).

## Discussion

SNHL can be caused by primary degeneration of SGNs or by secondary degeneration of these neurons after HC loss (44–46). Replacement and maintenance of SGNs would be an important step in any attempt to restore auditory function in patients with damaged sensory neurons or HCs (47,48). Successful replacement of lost or damaged SGNs is likely to result in improved clinical outcomes for cochlear implant recipients.

Cell therapy approaches based on the use of neural crest derived stem cells are promising for the regeneration of different peripheral nervous system tissue injuries. Among the neural crest derived stem cells conserved in several regions of the adult body, human dental pulp represents an easily accessible source to isolate stem cells with low invasiveness and in substantial quantities (15,49). The exploration and application of dental-related sources of human neural crest-derived stem cells (i.e., hDPSCs) to SNHL could therefore facilitate the development of a clinically viable, cell-based cell therapy for this neurosensory deficit. Although the auditory sensory neurons are derived from the otic placode and not from the neural crest like the sensory neurons described above, they share similar neurogenic genes and signaling pathways during their specification and differentiation (50,51). Considering the future potential applications of hDPSCs, the focus of the present study was to develop optimal culture conditions for their transdifferentiation into ONP and SGN-like cells via neurosphere-mediated and direct otic neuronal induction methods. During the time course of these two steps of *in vitro* differentiation procedures, we systematically screened factors and signals that can promote otic neuronal lineage pathways. Using qPCR and immunocytochemistry approaches, we tracked the emergence of OP, OV, ONP and SGN-like phenotypes by monitoring the expression of a panel of known lineage gene markers.

The main finding of the first step (i.e., neurosphere assay) of our stepwise differentiation procedure is that concomitant inhibition/activation of BMP signaling, along with TGF-β inhibition, was effective in promoting differentiation toward neural crest, otic/placodal and otic neuronal progenitor cell fates. This initial step led to the generation of otic neurospehres expressing Sox2/Pax2 that include otic neuroprogenitor cells which can differentiate into otic neurons. These finding are in agreement with previous reports concerning the effect of dual SMAD inhibition on promoting neural lineage of stem cells and hDPSCs (15,52–54). The expression of a large panel of neural markers also supports the ability of neural crest stem cell to adapt to the signaling cues of their environment and their stemness as previously suggested (55).

BMP4 is a member of the TGFβ family, and previous studies have shown that BMP4 is expressed early in the regions of otocysts, which correlates with the later development of SGNs and HCs (56,57). It has also been confirmed that BMP4 plays an important role for directing differentiation of inner ear stem cells, hESCs, IPSCs and bone marrow stem cells (BMSCs) toward a sensory neural fate *in vitro* (23,25,58,59). Similarly, we observed upregulation of ONP gene markers (*Sox2*, Neurod1) following BMP activation and TGF-β inhibition. The combined expression of such genes is crucial for proper otic neurogenesis and upregulation of these markers is sufficient to define a subpopulation of otic neuroprogenitors (23–25,27).

During embryogenesis, the otic placode invaginates into the underlying mesenchyme to form the otic vesicle. The ventral region contains otic neurosensory progenitors that give rise to sensory neurons, including SGNs. SHH is required for ventral patterning of the inner ear and acts synergistically with all-trans retinoic acid (ATRA) to facilitate otic sensory neuronal differentiation (26,36,39). In addition, during inner ear development, WNT co-operates with other pathways, particularly NOTCH and FGF, to specify the otic placode and its neurosensory derivatives (60). The interaction between SHH, RA and WNT produces a patterning gradient which induces target cells into otic placode neurosensory fate (39,61).

To further explore the role of these factors, we reasoned that similar *in vivo,* a timely directed modulation of these pathways could promote initial SGN lineage from hDPSCs under culture conditions. To test this hypothesis, we challenged the generated neurospheres with either ATRA/SHH/CHIR under NT3/BDNF exposition (paradigm 1) or with NT3/BDNF (Paradigm 2) from day 7 to day 21 *in vitro*. We found that both *in vitro* paradigms yielded the differentiation of hDPSC-derived ONP towards SGN-like cells. We assume that ONPs, which express Pax2 and Bmp7, are induced under paradigm 1 and lead to SGN-like cells expressing a panel of neuronal markers (NEUN, SOX2, TUJ1), confirming the otic modulatory effects of the molecules used in this paradigm (13,23,27,43). However, under paradigm 2, in addition to the increased relative expression of Pax2 and Bmp7, the ONPs cultured in this paradigm downregulate Dlx5 and upregulate TrkB, suggesting a strong commitment toward otic neuronal lineage. TrkB is a known receptor of BDNF and its expression is detected by embryonic day E12.5 in mice during SGN early specification (62,63). It is also well established that BDNF and NT3 are implicated in cochlear innervation during inner ear morphogenesis (64). The expression of TrkB and TrkC, which are regulated by neurotrophins at 13 DIV and later at 21 DIV, along with the expression of other known neuronal markers, supports the potential of paradigm 2 to generate SGN-like cells from hDPSCs.

Another distinction between the differentiating paradigms used in this study is related to prolongation of culture period beyond 21 DIV. Only, the cells differentiated under the paradigm 2 continued to growth with a substantial survival ratio by 32 DIV when compared to the cells differentiated under the paradigm 1. These observations could probably be related to the high number of GFAP immunopositive cells in culture paradigm 2. This subpopulation of differentiated GFAP immunopositive cells observed at 21 DIV may include GFAP+ cells that can be either Schwann cells precursors or glial cells, and would represent the cell population derived from the stepwise differentiation of hDPSCs (65,66). It is well known that glial and Schwann cells play an important role in the neurotrophic support and survival of SGN during inner ear development (67,68), which can explain the extended maturation of cell cultures of paradigm2 in contrast to cultures of paradigm1.

The SGN-like cells at 32 DIV are characterized by the co-expression of both SOX2 and BRN3A. This co-expression of these two known key markers, which was observed only in cells differentiated in paradigm 2, could indicate a possible early differentiation of some otic glutamatergic neurons (69). These SGN like-cells also express otic neuronal markers such as TRKC and PRPH, which corroborate the SGN phenotype reported by other *in vitro* models and the *in vivo* phenotype of SGN type 1 precursors, suggested by recent reports of scRNA seq analysis (69–71).

Additionally, we explored the nanomechanical properties of our generated SGN like-cell at 32 DIV differentiated in cultures of paradigm 2 and compared them to *in vivo* SGN equivalents and to undifferentiated hDPSCs. This nanomechanical characterization offers a high resolution cartography and meaningful information about the stiffness of biological samples (reviewed by (72)), allowing to distinguish between different cell populations including the neuronal cell types, and to conclude about similarities between their cytoskeletal compositions (73–75). Our explorations revealed that *in vitro* SGN-like cells and *in vivo* SGNs share the same Young’s modulus, which means that they have the same nanomechanical properties that are different from undifferentiated hDPSCs. Our initial explorations are correlated with a recent report that explored the optimal matrix mechanical properties to generate otic neurosensory progenitors, which is around 3 KPa (76), suggesting our SGN-like cells *in vitro* are similar to their in *vivo* SGN counterparts. Altogether, these data support the efficiency of our reliable protocol to generate SGN-like cells under culture conditions of the paradigm 2.

We also investigated the neuronal differentiation of hDPSCs-derived ONPs when co-cultured with rat postnatal SGN explants to evaluate the potential of using these cells to restore SGNs. Co-culture experiments offer an *in vitro* model to assess the therapeutic potential of human ONPs and *in vitro* neurons (31,77). When we cultured 13 DIV ONP with SGN explants, we observed preferential projections of afferent SGN explant neurons toward the ONPs. This suggests that ONPs might provide a supportive or trophic factor to SGNs that could offer therapeutic benefits in the context of preservation or regeneration of neuronal contacts from SGNs. As our hDPSCs-derived ONPs contains a subpopulation of glial and Schwann cells, we suggest that ONPs may share the same secretome features as the hDPSCs from which they derive. This hypothesis is related to the known pleiotropic secretome of hDPSCs which is very rich in neurotrophic and growth factors (NT3, IGF) that promote the survival and differentiation of neuronal cells (6,78). Furthermore, a recent report showed a similar effect of neuronal projections of SGNs due to secreted factors from otic pericytes in a co-culture system (79). Future studies are required to explore the nature of these contacts and the presence of synaptic markers, in addition to the electrophysiological properties of SGN like cells derived at 32 DIV, and whether trophic factors are secreted by ONPs, since the findings of the present study support a strong therapeutic potential of ONPs that needs to be uncovered.

In this study, we have extended the range of cell fate derivatives available for these hDPSCs to include SGN-like cells and we have demonstrated their nanomechanical and neuronal re-connection properties using co-cultures of human otic progenitors and SGN explants. Additional studies are required to establish the conditions required for their engraftment potential in animal models of auditory neuropathy, and such studies will be essential in terms of clinical use, which may be combined with cochlear implants to improve hearing.

## Ethical Approval

All experiments for human dental pup cells were performed in accordance with the local ethics committee (Comité de protection des Personnes, Centre Hospitalier de Montpellier)

## Acknowledgments

We thank the staff of the MRI-DBS-Optique facility (Elodie JUBLANC & Vicky DIAKOU) for the help with image acquisition and analysis and the RHEM-IGMM facility (Iria Gonzalez-Dopeso Reyes) for the help with immunochemistry. We also thank the staff of the MRI-IGMM facility (Stéphanie VIALA & Myriam Boyer-Clavel) for help with Cytometry experiments and the LabEx CEMEB for qPCR facilties. The graphical elements in Figure-S5 were designed and drawn by Zhanna Santybayeva (illustration4science.com).

## Limitation of the study

Although we were successful in deriving ONP and SGN-like cells from hDPSCs, some limitations of the study must be taken into consideration. While we were able to generate SGN-like cells expressing a comprehensive panel of otic neuronal markers and with the expected morphology, we did not assess the functionality of these cells, which should be investigated in future with electrophysiological studies. Moreover, cells in a more mature state of otic neuronal differentiation (i.e., 32 DIV) should be considered for future co-culture experiments to assess whether the level of maturation state influences neurite outgrowth and/or survival. Similarly, we did not investigate the effect of undifferentiated hDPSCs as a control for the effect of ONPs on neurite outgrowth in our co-cultures. The composition and effects of the culture medium from ONP could also be investigated and compared to conditioned medium from hDPSCs, which has previously been reported. Finally, we would like to obtain a through representative nanomechanical characterization of the SGNs generated in this study by analyzing SGNs harvested from human inner ear biopsies and SGNs from other *in vitro* differentiation models (i.e. iPSC and ESC-otic neuronal derivatives, and comparing them to our *in vitro* model system.

## List of nonstandard abbreviations

A-MEM: Alpha Modified Eagle Medium
BDNF: Brain-Derived Neurotrophic Factor
BMP: Bone Morphogenic Protein
BMSC: Bone marrow stem cells
DMEM/F12: Dulbecco’s Modified Eagle Medium: Nutrient Mixture F-12
DPBS: Dulbecco’s Phosphate Buffered Saline
FBS: Fetal Bovine Serum
hDPSC: Human Dental Pulp Stem Cell
HC: Hair Cell
MSC: Mesenchymal Stem Cell
NC: Neural Crest
OC: Organ of Corti
ONP: Otic Neuronal Progenitor
OP: Otic Placode
OV: Otic Vesicle
PBS: Phosphate Buffered Saline
PCR: Polymerase Chain Reactio
NT-3: Neurotrophin-3
RA: Retinoic Acid
SGN: Spiral Ganglion Neuron
SNHL: Sensorineural Hearing Loss
SV: Stria Vasculari

