## Supplemental data for "Transdifferentiation of Human Dental Pulp Mesenchymal Stem Cells into Spiral Ganglion-like Neurons"

### Supplemental Figure S1

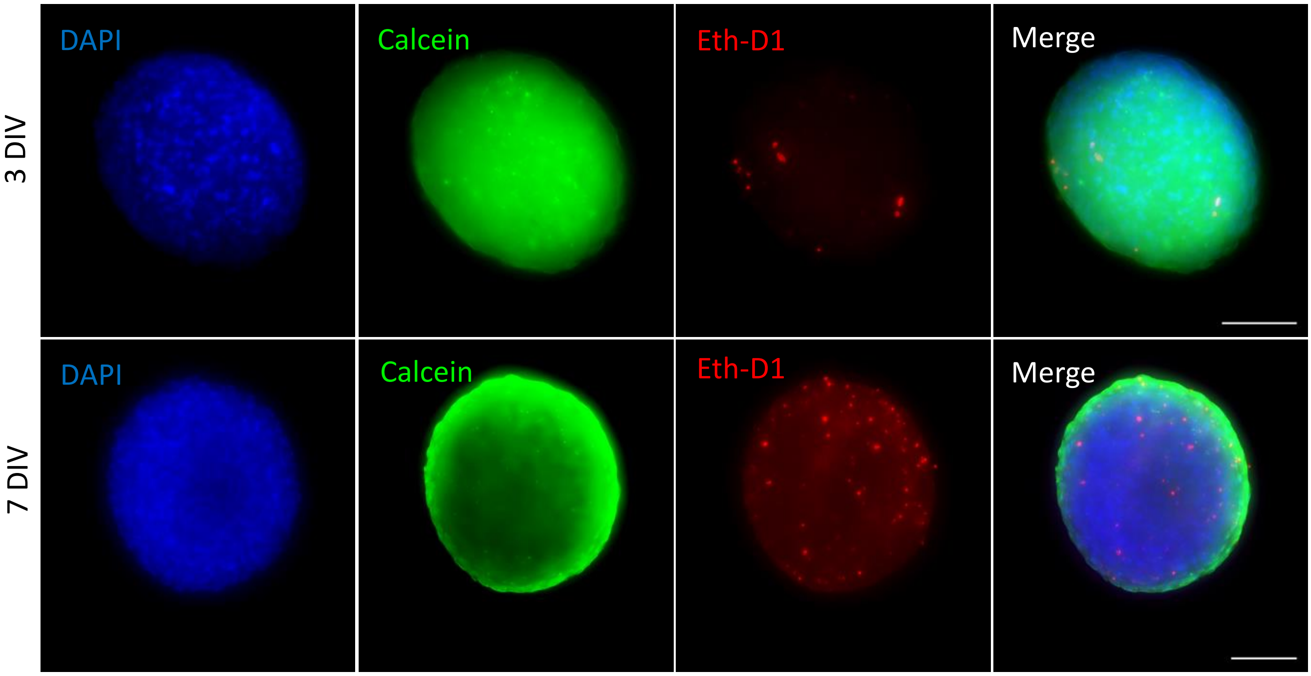


**Figure S1 -related to Figure 2. Representative fluorescent images from Live/dead assay on neurospheres at day 3 and day 7 *in vitro***.

The Live/dead staining distinguishes living from dead cells within neurospheres. Live cells (Calcein) were marked as green in the cytoplasm, and dead cells (EthD) were marked as red in the nuclei. The nuclei were counterstained with DAPI (shown in blue). Scale bar = 100 µm

### Supplemental Figure S2

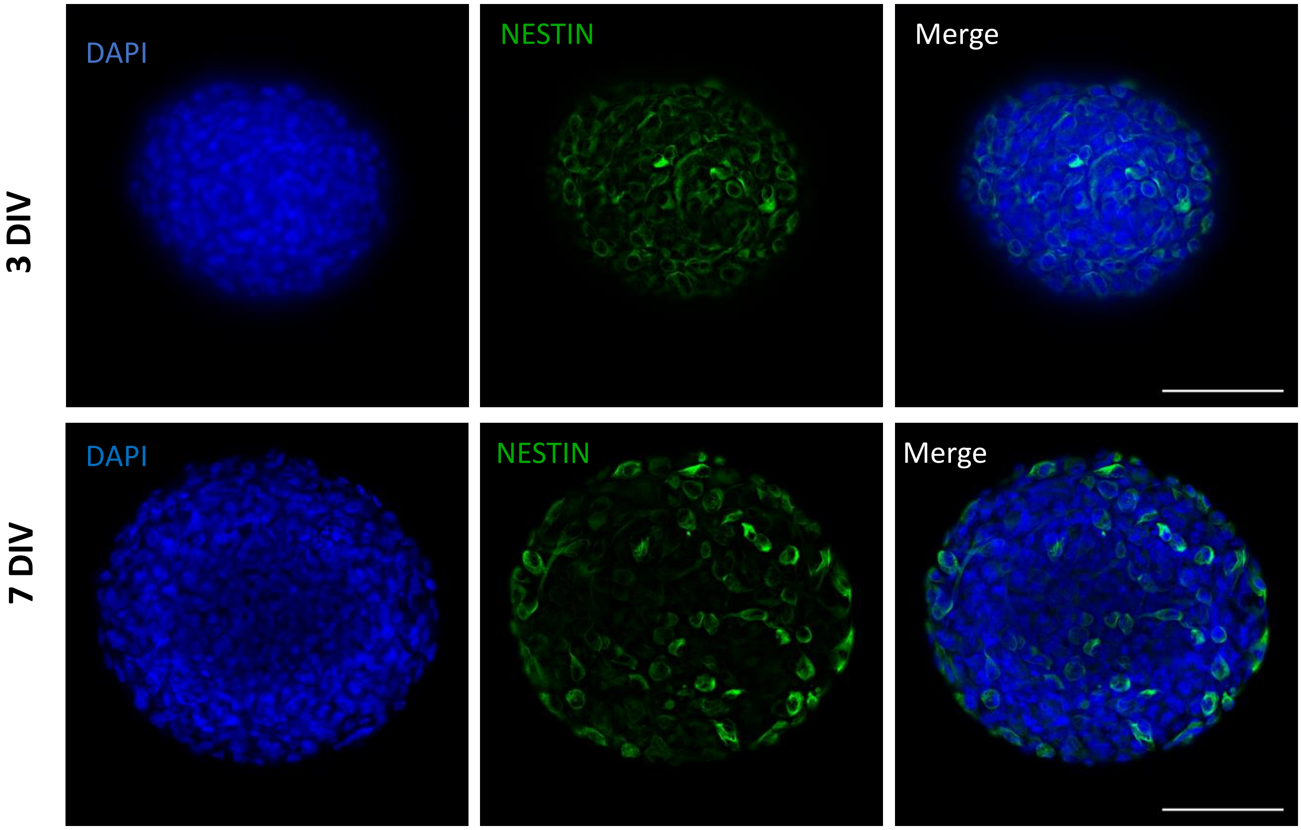


**Figure S2- related to Figure 3B.**

Representative images showing NESTIN (shown in green) immunopositive cells at day 3 at day 7 of differentiation of the protocol. Cell nuclei were counterstained with DAPI (blue). Scale bars = 100 µm.

### Supplemental Figure S3

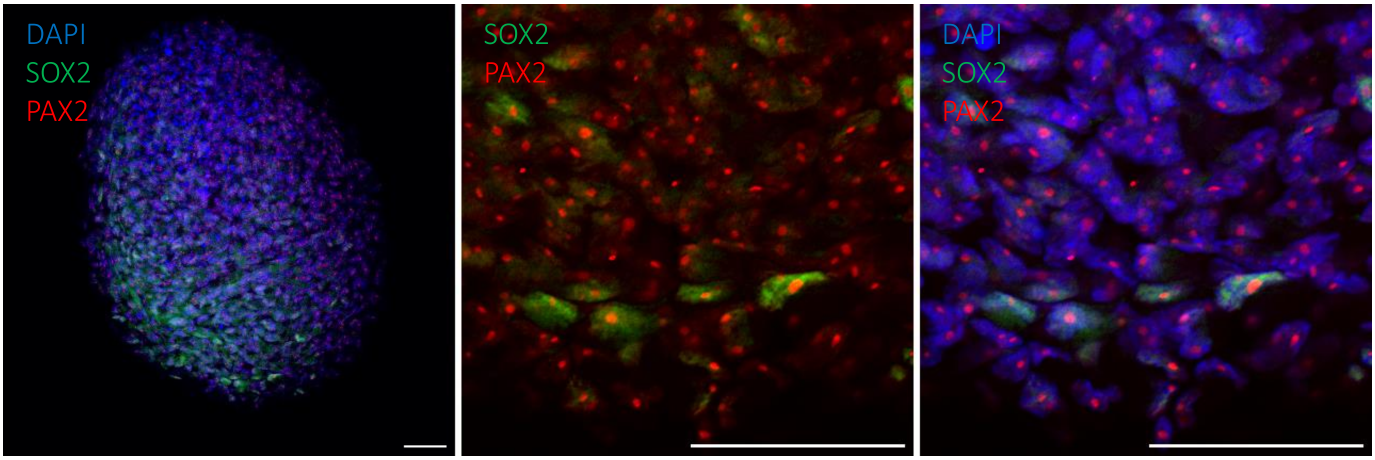


**Figure S3- related to Figure 3B.**

Representative images showing SOX2 (shown in green) and PAX2(shown in red) double immuno+ cells at 7 DIV . Cell nuclei were counterstained with DAPI (blue). Scale bars = 50 µm.

### Supplemental Figure S4

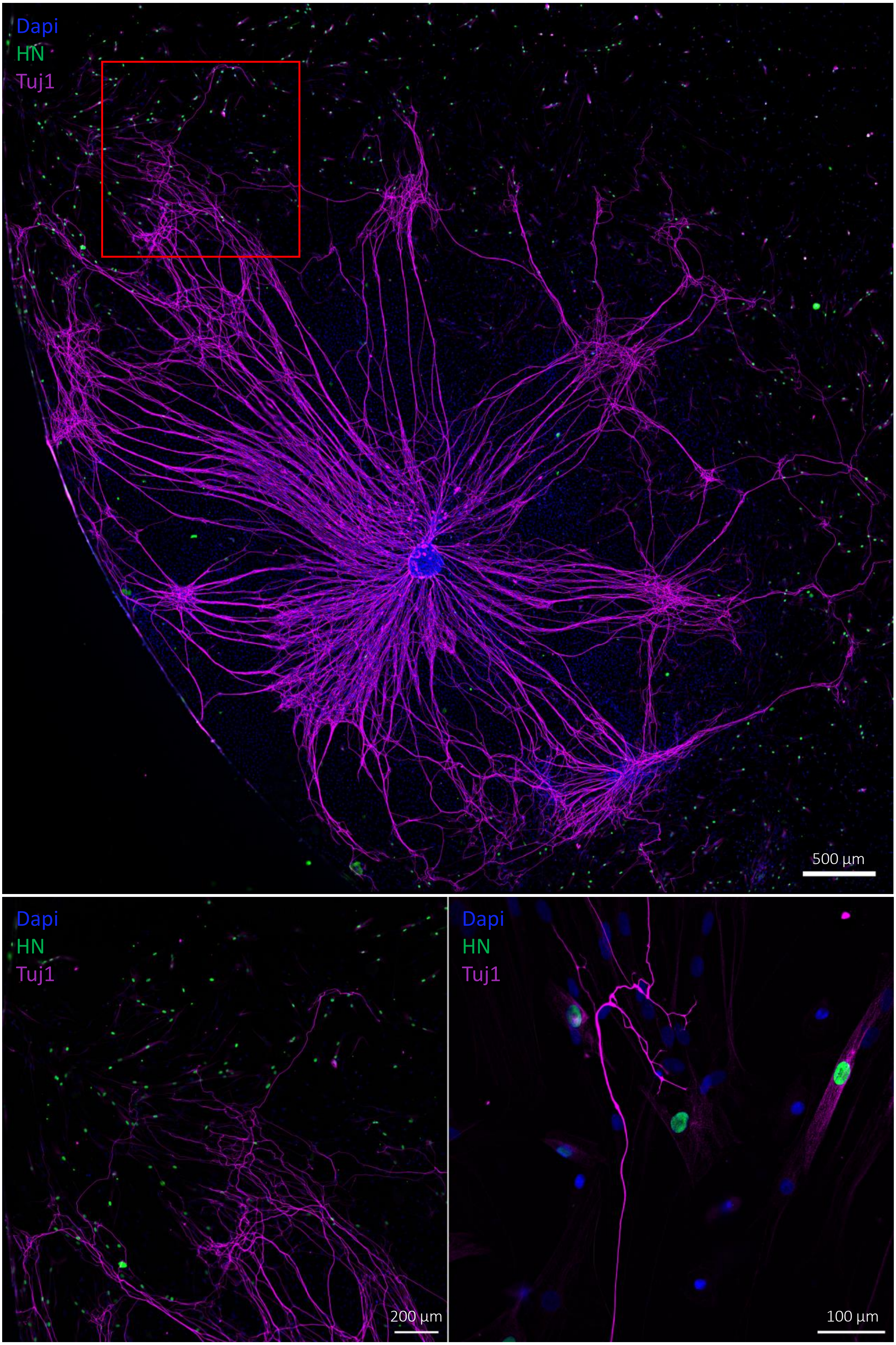

**Figure S4- related to Figure 10 .**

Representative fluorescent images of the neurite outgrowth toward the ONPS in co-culture. neurites (stained with Magenta= TUJ1) project toward the ONPs ( green= human nuclei & Magenta = TUJ1). Dapi was used to counterstain the nuclei.

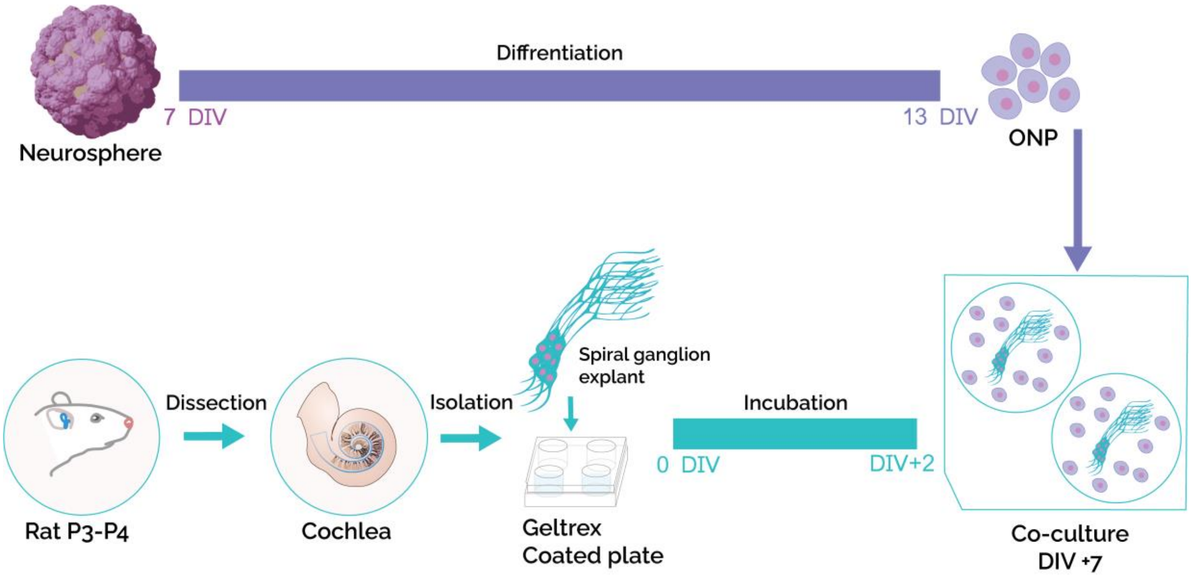


**Figure S5- schematic representation of the co-culture protocol.**

Neuroshperes were generated from hDPSCs and differentiated to ONP until 13 DIV. In parallel, Cochlear explants were dissected from P3 rats and SG was isolated and put in culture on Gletrex coated plates for 48h. ONPs were then detached using Accutase (Sigma) and co cultured with SGN explants for 5 additional days.

**TABLE S1.**

| Family | Gene | Forward | Reverse | Acces number |
|---|---|---|---|---|
| Neural Ectoderm | Ncad | TGGGAAATGGAAACTTGATGGC | AATCTGCAGGCTCACTGCTC | NM_001308176.2 |
| | Pax6 | TTGCCCGAGAAAGACTAGCA | TCTCCATTTGGCCCTTCGATTA | NM_001368892.2 |
| Non-Neural Ectoderm | Eya1 | ACAGCCGACGGGTCTTTAA | TTGGTCGTGGGCTGAAACTA | NM_172058.4 |
| | Six1 | GGTTTAAGAACCGGAGGCAAA | TGCTTGTTGGAGGAGGAGTTA | NM_005982.4 |
| | Dlx5 | GCTAGCTCCTACCACCAGTAC | GGTTTGCCATTCACCATTCTCA | NM_005221.6 |
| | Tfap | TTTCAGCCATGGACCGTCA | GGGAGATTGACCTACAGTGC | NM_001032280.3 |
| | Foxi1 | GACAAGCGCCTCACTCTCA | CCGGCCTTGCTCTTGTTGTA | NM_144769.4 |
| | Pax2 | CGGCTGTGTCAGCAAAATCC | GCTTGGAGCCACCGATCA | NM_000278.5 |
| | Pax8 | GCCCAGTGTCAGCTCCATTA | GCTGTCCATAGGGAGGTTGAA | NM_013992.4 |
| | Ecad | CAGGAGTCATCAGTGTGGTCA | CAAAATCCAAGCCCGTGGTG | NM_001317184.2 |
| | Lhx2 | CAAAAGACGGGCCTCACCAA | CGTAAGAGGTTGCGCCTGAA | XM_006717323.3 |
| | Bmp7 | ACGTTCCGGATCAGCGTTTA | CTGTCGAGCAGGAAGAGATCC | NM_001719.3 |
| | Dnmt3a | GAGCGGGTTGTGAGAAGGAA | TCCTGCAATGACCTTGGCTT | NM_153759.3 |
| Neural Crest | Foxd3 | CTCATGGCCACCCACCAA | GGAGAGTGGCACGCTAAGAA | NM_012183.3 |
| | Sox10 | TCGCGGACCAGTACCC | GCGCTTGTCACTTTCGTTCA | NM_006941.4 |
| | Snail1 | CGAGTGGTTCTTCTGCGCTA | GGGCTGCTGGAAGGTAAACT | NM_005985.4 |
| | Sox2 | AGCTCGCAGACCTACATGAA | GGAGTGGGAGGAAGAGGTAAC | NM_003106.4 |
| | Plp1 | TCCACCCTCAATCCACATTTC | TGGCTAGTCTGCTTTGTGGC | NM_001128834 |

| | | | | |
|---|---|---|---|---|
| | | C | | .3 |
| | Nestin | GTGGCTCCAAGACTTCCCTC | GGTGTCTCAAGGGTAGCAGG | NM_006617.2 |
| | Zic1 | GCGCGCTCCGAGAATTTA | CCCTCAAACTCGCACTTGAA | NM_003412.4 |
| | Pax3 | GCGGTCTGTGATCGAAACA | TCCTCCTCTTCACCTTTCCC | NM_181461.4 |
| | Pax7 | ACAGCATCGACGGCATCC | CAGGTTCCGACTCCACATCC | NM_013945.3 |
| SGN | Ascl1 | CGGTCTCATCCTACTCGTCG | CGCCACTGACAAGAAAGCAC | NM_004316.4 |
| | Neurod1 | TGACTGATTGCACCAGCCCT | TTCTCAAACTCGGCGGACGG | NM_002500.5 |
| | Neurog1 | AGCGCCTTTCTATCTGTCCG | AGGAAGCCGGATAGGTCACT | NM_006161.3 |
| | Neurog2 | AGGCCAAAGTCACAGCAACG | CCAAGGTCTCGGATTTGACG | |
| | Dll1 | CTCAGGGGAGGAGAAGGGG | GAGAAACGGGAGTCTTGCCA | NM_005618.4 |
| | Brn3a | CTGAGCACAAGTACCCGTCG | GCTTGAAAGGATGGCTCTTGC | NM_006237.4 |
| | Tlx3 | GTTCCAAAACCGGAGGACCA | CTGGATGGAGTCGTTGAGGC | NM_021025.4 |
| | Pou3f4 | CCCATTTCGGTTCACCTCCA | GCAGAGAATGCCTATCCCCC | NM_000307.5 |
| | Tbx1 | TCGACAAGCTCAAGCTGAC | GCTGGTATCTGTGCATGGAA | NM_080646.2 |
| | Nfl | ACAAGCAGAACGCCGACATC | GGTCTCCTCGCCTTCCAAGA | NM_006158.5 |
| | Nfm | TCCGGCAGTGATCGGAAGAG | AGCCATTTCCCACTTTGTGC | |
| | Nfh | CCGACATTGCCTCCTACCAG | GCCATCTCCCACTTGGTGTT | NM_021076.4 |
| | Notch1(receptor) | ACGGCGTGAACACCTACAA | TGGCACTCGTCCACATCC | XM_011518717.2 |
| | JAG1 (ligand) | AACAAAGGCTTCACGGGAAC | CAAGTGCCACCGTTTCTACAA | NM_000214.3 |
| | Isl1 | TCGCCTTGCAGAGTGACATA | CCCGGTCCTCCTTCTGAAAA | NM_002202.3 |
| | Peripherin | GCCGGAAGACGGTTCTGAT | TAGGGTTTGGGCTTTGAGCA | NM_006262.4 |
| | Trkb | GGAATTGGGTTGGAGCAGGA | GGGGCGCAGATTCCTTGTTA | |
| | Trkc | CACCCCTTCCTGATGTGGAC | GCCATTGTCCTCACTCGTCA | |
| | Vglut1 | AGGAGCGCAAGTACATCGAG | CGCCAGGGAGTGCTAAACTT | |
| | Bdnf | AGCCTTTTCCTCCTGCTGTG | GCAGCCTTCATGCAACCAAA | NM_001143805 |

| | | | | |
|---|---|---|---|---|
| | | | | .1 |
| | Nt3 | CGCACATCTGGGACCCCT | TGGACATCACCTTGTTCACCT | XM_011520963.2 |
| Schwann Cells | P75 | CAGGACAAGCAGAACACCGT | GGTGTGGACCGTGTAATCCA | NM_002507.4 |
| | Ncam | GATGCGACCATCCACCTCAA | TCTCTGGTCGAGTCCACGAA | |
| | S100 | AGGAGCTGAAAGAGCTGCTG | TGTCCACAGCATCCACATCC | NM_006271.2 |
| | Oct6 | TGGACTCTTTTTGTTCGGTTGC | CGTCCGGGTGTTTGGTTTTG | NM_002699.4 |
| Endoderm | Cxcr4 | CCCGACTTCATCTTTGCCAAC | ACACAACCACCCACAAGTCA | NM_003467.3 |
| | Sox17 | CACAACGCCGAGTTGAGCAA | GCTCTGCCTCCTCCACGAA | NM_022454.4 |
| | Gata4 | AAAACGGAAGCCCAAGAACC | AAGGCTCTCACTGCCTGAA | NM_001308094.2 |
| | Gata6 | GGGCTCTACAGCAAGATGAAC | GTTGGCACAGGACAATCCAA | NM_005257.6 |
| | Epcam | GTGCTGGTGTGTGAACACTG | GAAGTGCAGTCCGCAAACTT | NM_002354.3 |
| | FoxA2 | ACTGGAGCAGCTACTATGCA | TGTTCATGCCGTTCATCCC | NM_021784.5 |
| Mesoderm | Bra/T | CGCTTCAAGGAGCTCACCAA | GCCAGACACGTTCACCTTCA | NM_003181.4 |
| | Hand1 | CAAGCGGAAAAGGGAGCTG | CAGCCGGTGCGTCCTTTAAT | |
| | Sox7 | GGCCAAGGACGAGAGGAAAC | TCCGCCTCGTCCACGTA | NM_031439.4 |
| | Mixl1 | GTACCCCGACATCCACTTGC | ACCTGGAAGAGGGGAGAAAATAA | |
| Housekeeping | Rps18 | CCGCCATGTCTCTAGTGATCC | GGTGAGGTCGATGTCTGCTT | NM_011296.2 |

TABLE S2.

| Antigen | Host | Brand | Ref | Dillution |
|---|---|---|---|---|
| SOX2 | Mouse | Abcam | ab97959 | 1:100 |
| SOX2 | Rabbit | Merck | AB5603 | 1:200 |
| STRO1 | Mouse | Invitrogen | 398401 | 1:150 |
| NESTIN | Mouse | Abcam | ab18102 | 1:250 |
| PAX2 | Rabbit | Abcam | ab79389 | 1:200 |
| B3-TUBILIN | Mouse | Abcam | ab78078 | 1:250 |
| MAP2 | Mouse | Invitrogen | 13-1500 | 1:200 |
| TRKc | Rabbit | Cell signaling | C44H5 | 1:100 |
| NEUN | Rabbit | Cell signaling | D3S3I | 1:100 |
| PERIPHERIN | Rabbit | Abcam | ab4666 | 1:250 |
| NF-M | Mouse | DSHB | 2H3-C | 1:200 |
| GFAP | Mouse | DSHB | 8-1E7-S | 1:150 |
